# The bacterial amyloids phenol soluble modulins from *Staphylococcus aureus* catalyze alpha-synuclein aggregation

**DOI:** 10.1101/2021.03.10.434765

**Authors:** Caroline Haikal, Lei Ortigosa, Katja Bernfur, Alexander Svanbergsson, Sara Linse, Jia-Yi Li

## Abstract

Aggregated α-synuclein (α-syn) is the main constituent of Lewy bodies, the main pathological hallmark of Parkinson’s disease (PD). Environmental factors are thought to be potential triggers capable of initiating the aggregation of the otherwise monomeric α-syn. Braak’s seminal work redirected attention to the intestine and recent reports of dysbiosis have highlighted the potential causative role that the microbiome might play in the pathology of PD. *Staphylococcus aureus* is a bacterium carried by 30-70% of the general population. It has been shown to produce functional amyloids, called Phenol Soluble Modulins (PSMαs). Here, we studied the kinetics of α-syn aggregation under quiescent conditions in the presence or absence of four different PSMα peptides and observed a remarkable shortening of the lag phase in their presence. Whereas pure α-syn monomer did not aggregate up to 450 h after initiation of the experiment in neither neutral nor mildly acidic buffer, the addition of different PSMα peptides resulted in an almost immediate increase in the Thioflavin T (ThT) fluorescence. Despite similar peptide sequences, the different PSMα peptides displayed distinct effects on the kinetics of α-syn aggregation. Kinetic analyses of the data suggest that while all four peptides catalyze α-syn aggregation, the underlying mechanisms might differ with a model of nucleation and elongation fitting the α-syn aggregation induced by PSMα2 but not PSMα1. The results of immunogold TEM imply that the aggregates are fibrillar and composed of α-syn. Addition of the co-aggregated materials to HEK cells expressing the A53T α-syn variant fused to GFP was found to catalyze α-syn aggregation and phosphorylation in the cells. Our results provide evidence of a potential trigger of synucleinopathies and could have implications for the prevention of the diseases.

## Introduction

Aggregated alpha-synuclein (α-syn) is a pathological hallmark of Parkinson’s disease (PD), Multiple System Atrophy and Lewy Body Dementia. The majority of synucleinopathies are of idiopathic nature and the environment is thought to play a causative role. In Braak’s landmark studies, α-syn inclusions were identified in the dorsal motor nucleus of the vagus at very early stages of PD. It was hypothesized that PD pathology could be initiated peripherally, either in the gastrointestinal or olfactory systems, wherefrom it would spread to the brain (1, 2). Several studies in rodents have shown that α-syn can propagate from the intestinal lumen, wall or enteric neurons to the brain(*3–6*). Other studies have shown propagation of preformed α-syn fibrils from the olfactory bulb to the amygdala and enterohinal cortex (*7–9*). For a deeper understanding of PD and other synucleinopathies, it is as such of interest to examine peripheral factors which could trigger the aggregation of the otherwise soluble α-syn.

Both the olfactory cavities and the gastrointestinal tract harbor large numbers of microorganisms that are known to prime and modulate the immune system as well as directly modulate α-syn aggregation. Both LPS and bacterial chaperones have been shown to affect the kinetics of aggregation and toxicity of α-syn (*10–12*). Cross-seeding of α-syn has previously been demonstrated not only with human amyloid proteins (tau) (*13*) and Aβ42 (*14*) but also with amyloid proteins of bacterial origin (Curli from E. coli) (*11, 15*). Several bacteria and fungi produce amyloids, which often help in the maintenance of a biofilm and adherence (*16*). One such bacterium is the gram-positive *Staphylococcus aureus*.

*S. aureus* is the bacterium responsible for the majority of skin and soft-tissue infections (*17*) as well as a common skin, nose and even GI commensal (*18–21*). *S. aureus* produces amyloid proteins called phenol soluble modulins (PSMs). The α type of the PSMs comprise four different hydrophobic 20-25 amino-acid long peptides, often formylated at the N-terminal. The production of PSMα peptides by *S. aureus* is stringently regulated and is associated with virulence. PSMα peptides are secreted by *S. aureus* at high concentrations, with high nanomolar concentrations acting as chemoattractants for neutrophils and micromolar concentrations causing cytolysis (*22, 23*). Their expression has been shown to be upregulated upon *S. aureus* phagocytosis by neutrophils promoting phagosome lysis and bacterial escape (*24, 25*). PSMα peptides can rapidly induce neutrophil extracellular trap (NET) formation (*23*). Interestingly, NETs have been shown to colocalize with amyloids in human tissues (*26*).

PSMα peptides produced by *S. aureus* have been shown to directly activate sensory neurons of the skin and increase firing in cultured DRG neurons by disrupting membranes and forming pores, allowing cation influx and depolarization (*27, 28*). In contrast, they have been shown to inhibit the firing of sensory neurons in the enteric nervous system leading to modulation of the secretion and motility of the intestine (*29*). Nociceptive hypersensitivity and GI disturbances have been described both in PD patients and PD rodent models. Skin biopsies have revealed small fiber neuropathies or mixed fiber polyneuropathy in PD patients (*30*) and Braak has described α-syn deposits in neurons of the lamina I of the spinal cord, which directly project to the thalamus from peripheral nociceptive neurons (*31*). Lewy pathology has also been described in the ENS (*32*) and neurons of the DRG (*33*). PSMα peptides could as such interact with α-syn in peripheral sensory neurons.

In the present study, we hypothesized that these PSMα peptides could modulate α-syn aggregation. We monitored the aggregation kinetics of α-syn *in vitro* upon the addition of different PSMαs at a range of concentrations. The PSMαs induced rapid aggregation of monomeric α-syn into fibrils, which could induce seeding and phosphorylation in HEK cells expressing the A53T variant of α-syn fused to GFP.

## Materials and Methods

### α-Syn production and purification

Human wildtype α-syn was expressed and purified using heat treatment, ion exchange and gel filtration chromatography as previously described (*34*). α-Syn samples were aliquoted and stored at −20 °C.

### PSMα production

The four different PSMα peptides were produced by chemical synthesis and isolated using Reversed Phase HPLC to 95% purity (synthesis and purification purchased from EMC microcollections GmbH as lyophilized peptides) with the following sequences:

PSMα1: Formyl-MGIIAGIIKVIKSLIEQFTGK
PSMα2: Formyl-MGIIAGIIKFIKGLIEKFTGK
PSMα3: Formyl-MEFVAKLFKFFKDLLGKFLGNN
PSMα4: Formyl-MAIVGTIIKIIKAIIDIFAK

The lyophilized peptides were stored at − 80 °C and used without further purification in all experiments except one. For one experiment, PSMα2 was further purified using size exclusion chromatography. The peptide was dissolved at 1.25 mg/ml in 6 M GuHCl, 10 mM MES and purified using a Superdex Peptide 10/300 gl column in 10 mM Tris, 50 mM NaCl, pH 7.6, with absorbance measurements at 214 nm and 256 nm.

### Aggregation kinetics experiments

In order to obtain reproducible data, monomeric α-syn was isolated immediately prior to each experiment. Purified α-syn was lyophilized, dissolved in 6 M GuHCl,10 mM MES, pH 5.5 and purified using size exclusion chromatography in 10 mM MES, pH 5.5 or 10 mM Tris, 50 mM NaCl, pH 7.6 at the start of every experiment. Buffer solutions were filtered and degassed prior to each run. The purification was monitored by UV absorbance at 280 nm and only the central monomer peak collected. The absorbance of α-syn was measured at 280 nm and an extinction coefficient of 5960 M^−1^ cm^−1^ was used to calculate the concentration. Purified α-syn monomer was kept on ice to prevent aggregation.

The PSMα peptides were dissolved in DMSO to a concentration of 2 mg/ml and then diluted in buffer. The final DMSO concentration ranged from 8.9 to 0.1% v/v. Complementary α-syn aggregation experiments without PSMα peptides were performed with the same DMSO concentrations.

To follow the aggregation process, samples were aliquoted into 96-well black Corning polystyrene half-area microtiter plates with a non-binding surface (Corning 3881) in the presence of 20 μM ThT and 0.01% NaN_3_. Replicates without ThT were also prepared for additions to cell cultures. For the first round of experiments, the α-syn concentration was kept constant at 25 μM and PSMαs added at 5 different final concentrations: 77.7, 25.9, 8.6, 2.9, and 0.97 μM. Plates were incubated under quiescent conditions at 37 °C and the fluorescence was measured for up to 400 h (excitation filter 440 nm and emission filter 480 nm). Each experiment was repeated 1-3 times. Data from one experiment, representative of all, is shown in the results section.

For the second round of experiments, the PSMα concentrations were kept constant at 8.6 μM and α-syn added at concentrations of 116, 77.7, 25.9, 8.6, 2.9, and 0.97 μM. These experiments were performed in 10 mM Tris 50 mM NaCl pH 7.6 in the presence of 20 μM ThT and 0.01% NaN_3_.

### Kinetic analysis

The data sets from each experiments were analyzed using Amylofit (*35*). The ThT curves were normalized and the half time, defined as the point of time at which the ThT fluorescence has reached half-way in between the starting baseline and ending plateau, was extracted. The averages and standard deviations of the halftimes of 3 or 4 replicates for each treatment condition were plotted against the concentrations of PSMα peptides or α-syn. A model of nucleation and elongation was fitted to the kinetic data from the aggregation of α-syn in the presence of varying concentrations of PSMα1 and PSMα2 peptides.

### Gel and Mass Spectrometry

The samples were analyzed by SDS-PAGE using Novex 10-20% Tricine precast gels and Tricine SDS Running buffer. 20 μl of each sample were mixed with a five times concentrated gel loading buffer (4 ml 4.5 M Tris pH = 8.45; 4.8 ml glycerol; 1 g SDS; 1 ml ß-mercaptoethanol; 1 ml 1 % Coomassie Blue). 10 μl of each mixture were loaded, and the gel was run at 70 mV for 15 minutes, followed by 120 mV for 1 hour. The gel was then stained with InstantBlue™ overnight. Gel bands of interest were excised and subjected to in-gel digestion. Gel pieces (1×1 mm) were washed twice in 50 mM ammonium bicarbonate (NH_4_HCO_3_)/50% acetonitrile (ACN) followed by dehydrated in 100% ACN. Digestion was performed by adding 50 mM NH_4_HCO_3_ with 12 ng/μl sequencing-grade modified trypsin (Promega, Madison, WI, USA), incubation on ice for 4 h before overnight incubation at 37 °C. The next day Trifluoroacetic acid (TFA) was added to a final concentration of 0.5% and the peptide containing solution above the gel pieces was withdrawn and used for further analysis. Mass spectrometry analysis was performed in reflector positive mode on an Autoflex Speed MALDI TOF/TOF mass spectrometer (Bruker Daltonics, Bremen, Germany). All peptide mass spectra were externally calibrated using Bruker Peptide Calibration Standard II.

### Electron microscopy

The protein samples were examined under TEM. Samples were diluted 1:500 in sterile PBS (HyClone 10126473) and 5 *μ*l diluted sample added onto carbon-coated grids (made in-house). After washing in dH2O, nonspecific binding was blocked with BSA (SigmaA2153) and the specimens then incubated with1:200 syn211 (sc-12767) for 1 h. Samples were again washed in dH2O, incubated with 1:20 10 nm gold-anti-mouse antibody (Ted Pella 15751), washed, fixed first with gluteraldehyde (Ted Pella 18427) and then uranyl acetate (Agar Sientific R1260A).

### HEK 293T culture

HEK 293T cells stably transduced with the A53T mutant of α-syn fused to GFP and were maintained in DMEM supplemented with 10% FBS and 1% P/S and split every three days.

For live-imaging experiments, cells were split 1:20,000 into black, clear bottom 96-well plates. Samples of α-syn and PSMα incubated separately or together were sonicated at 100% amplitude for 3 min (1 s on, 1 s off). 10 μl of each sample was added in 4 or 3 replicates. The plates were imaged for 48 h with 1-h intervals using a Nikon Ti-E microscope with 95% humidity 5% CO_2_.

### Phospho-α-syn staining of HEK 293T cells

After 48 h, the cells were fixed with phosphate-buffered paraformaldehyde pH 7.4 by slow addition into the cell medium to a final concentration of 2% for 20 minutes. The cells were then washed and kept at 4°C. Cells were then permeabilized and blocked with 5% NDS 0.25% Tween20 in PBS. Primary antibody pS129 α-syn (Abcam 51253) was added at 1:1000 overnight and the plates kept at 4°C. The cells were washed and incubated with 1:600 Cy3 conjugated anti-rabbit antibody (Jackson immunolabs 711-165-152), washed and then imaged again. For each well, 25 (5×5 square) images were acquired with a 20x objective.

### Quantification of aggregates

Quantification of GFP-aggregates and pS129 α-syn inclusions was performed using Cell profiler. In brief, aggregates and cells were segmented by diameter sizes. An Otsu adaptive thresholding strategy was used to identify aggregates whereas a global threshold was used for nuclei. The number of aggregates and cells were quantified per well. The number of aggregates and phosphorylated aggregates per 100 cells were determined per well and averages from 4 replicates plotted for each treatment condition.

## Results

### Phenol soluble modulins catalyze α-syn aggregation

We hypothesized that the PSMαs could induce α-syn aggregation at physiologically relevant concentrations of 77.7 to 0.97 μM. To study the effects the kinetics under conditions of primary and secondary nucleation (pH 5.5 and low ionic strength), or only primary nucleation (pH 7.5 and moderate ionic strength), the experiments were repeated in two different buffers.

We first monitored the aggregation kinetics of 25 μM α-syn under quiescent conditions in the presence or absence of different PSMα peptides in mildly acidic MES buffer pH 5.5 with no salt. Under these conditions, all four types of PSMα significantly shortened the lag phase for α-syn aggregation to less than 20 h, and the ThT curves plateaued before 100 h even at low μM PSMα concentrations (Supplementary Figure1).

As the PSMα peptides would most likely interact with α-syn in the cytoplasm of sensory neurons or in the extracellular space, we wanted to examine the effect of the PSMαs on α-syn aggregation at neutral pH. Therefore, we repeated the experiments in 10 mM Tris, 50 mM NaCl, pH 7.6. Similarly, at neutral pH and moderate *I*, the addition of PSMα peptides induced a rapid increase in ThT fluorescence, which was not observed for α-syn incubated on its own or with DMSO at the same concentrations as used with the peptides (Figure 1, Supplementary Figure 2). At the examined concentrations, moderately faster aggregation was observed at mildly acidic pH at low *I*. Additionally, at low peptide concentrations, PSMα3 accelerated α-syn aggregation only at slightly acidic pH. The results indicate that PSMα peptides potently catalyze α-syn aggregation in buffers mimicking physiological conditions, rapidly increasing the ThT fluorescence even at low peptide concentrations.

**Figure 1.**
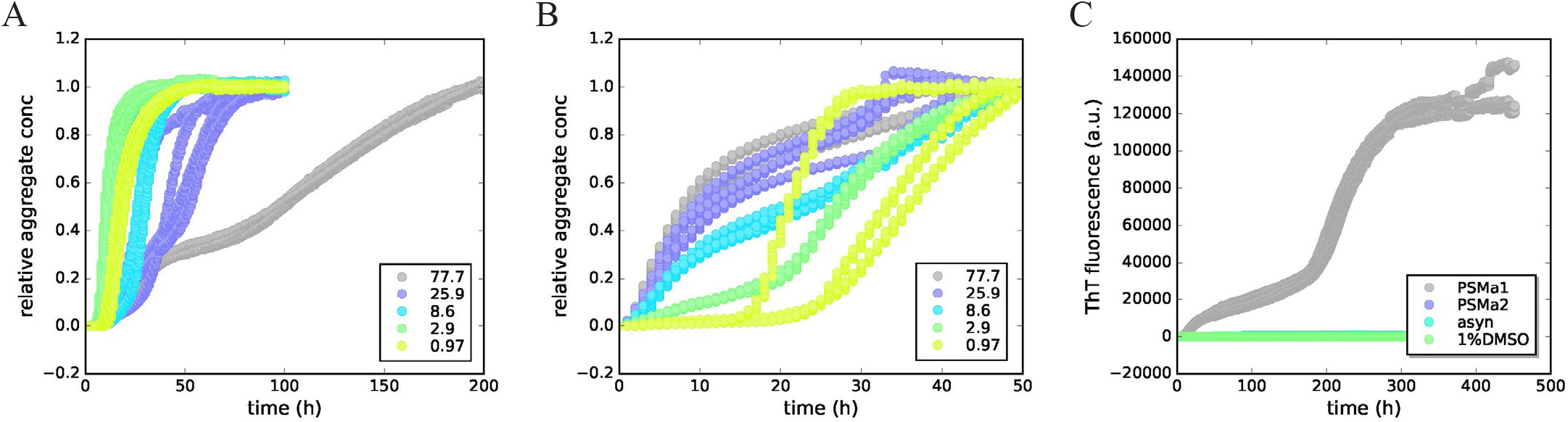
PSMα peptides induce rapid aggregation of α-syn. A. The normalized ThT fluorescence intensities reveal rapid aggregate formation upon addition of varying concentrations (μM) of A. PSMα1 and B. PSMα2 to 25 μM α-syn. C. Pure monomeric α-syn or α-syn supplemented with 1% DMSO did not aggregate under these conditions. 77.7 μM PSMα1 showed a stepwise increase in ThT fluorescence whereas PSMα2 at the same concentration did not. All experiments were performed in 10mM Tris with 50 mM NaCl, pH 7.6 at 37°C under quiescent conditions.

### DMSO does not affect PSMα modulation of α-syn aggregation kinetics

As the PSMα peptides were dissolved in DMSO, the samples with the peptide also contained DMSO, the concentration of which increased with the peptide concentration. To exclude the possibility that DMSO underlay the modulation of α-syn aggregation by the PSMαs, observed above, we added PSMα2 peptide to α-syn at 5 different concentrations and supplemented with DMSO to correspond to the highest DMSO concentration (8.9% v/v, equal to 1.25 M) in the previous experiments. The kinetic curves of PSMα2 dilutions with additional DMSO added to α-syn show similar trends as the PSMα2 dilutions without added DMSO (Supplementary Figure 3). Additionally, to further ensure the effects on α-syn aggregation observed were due to the peptides, and not DMSO or impurities, PSMα2 was purified with size exclusion chromatography. Different fractions of the central peak were added to freshly purified α-syn monomer in 10 mM Tris, 50 mM NaCl, pH 7.6 and compared to PSMα2 dissolved in DMSO. The results indicate similar kinetics for the purified PSMα2 fractions without DMSO to the original DMSO-dissolved peptide (Supplementary Figure 4).

### PSMα peptides induce α-syn fibril formation

To examine whether the observed ThT fluorescence increases report on α-syn aggregation, we imaged the samples by Transmission Electron Microscopy (TEM). Aggregates formed from α-syn in the presence or absence of PSMαs were detected by immunogold using anti-α-syn antibodies in the form of fibrils of approximately 10-20 nm in diameter (Figure 3, Supplementary Figure 5). α-syn incubated on its own did occasionally show individual fibrils despite the lack of increase in ThT signal in the kinetics experiments, in agreement with the formation of significant amounts of fibrils during the lag phase, although their concentration needs to reach about 1% of the final value to be detected by ThT bulk assay (*36, 37*). α-syn incubated on its own did, however, show higher background immunogold labeling, indicating high monomeric α-syn content.

### Different PSMαs modulate α-syn aggregation differently

Next, we wanted to elucidate the mechanisms underlying the PSMα-catalyzed α-syn aggregation. As a first step, we examined the effects of the different PSMα concentrations on the rate of α-syn aggregation. The time at which ThT intensity reached half the intensity of the plateau was calculated for each concentration of the four different PSMα peptides and plotted against the concentration of the peptides (Supplementary Figure 6). At pH 7.6, the double logarithmic plots of PSMα1 and PSMα4 show graphs with discontinuity indicating an optimal catalytic concentration above and below, which the lag phase is prolonged. In contrast, PSMα2 and PSMα3 exhibit linear curves with negative slopes, indicating a shortening of half-times as a function of concentration. This is interesting as PSMα1 and PSMα4 have previously been described to form fibrillar amyloid structures when aggregated on their own, as opposed to PSMα2 and PSMα3 which form α-helix-rich amorphous aggregates (*38, 39*). Indeed, in our hands, PSMα1 and PSMα4 regularly showed increases in ThT fluorescence, whereas PSMα2 and PSMα3 did not (Supplementary Figure 2). As such, at higher peptide concentrations, the aggregation of PSMα1 and PSMα4 could potentially compete with α-syn aggregation. A similar retardation of α-syn aggregation can be seen upon the addition of very high concentrations of lipid vesicles (*40*). This is thought to be due to the adsorption of α-syn monomer to the membrane, effectively lowering the free monomeric α-syn concentration. Thus, the smaller catalytic effect of α-syn aggregation in the presence of 77.7 μM PSMα1 or PSMα4 but not PSMα2 nor PSMα3 could indicate differential affinities for α-syn by the different PSMα peptides, or the presence of clusters of some of the peptides in solution.

The data for PSMα1 and PSMα2-induced α-syn aggregation were individually fitted to a model of nucleation and elongation (Supplementary Figure 7). The model fits the data of PSMα2 well, indicating PSMα2 could induce α-syn aggregation by primary heterogeneous nucleation. Clearly, for PSMα1, the model does not fit the data well. Interestingly, PSMα1 has been shown to aggregate primarily by secondary nucleation at neutral pH (*41*). To our knowledge, secondary nucleation has not been demonstrated for α-syn at neutral pH. Further studies are planned to elucidate the mechanisms underlying PSMα1-induced α-syn aggregation.

### PSMα-peptides induce α-syn aggregation by heterogeneous primary nucleation

To examine the α-syn-monomer-concentration-dependence of PSMα-induced aggregation the peptide concentration was kept constant and α-syn monomer concentration varied. For all peptides examined, a significant acceleration of aggregation of α-syn can again be observed (Figure 2, Supplementary Figure 8) compared to α-syn without the peptide, which did not show an increase of ThT signal. For the pure peptides, only PSMα1 can be seen to emit a ThT signal at 8.6 μM. The fluorescence intensity of this signal is similar to that emitted in the presence of low α-syn concentrations, yet low α-syn concentrations seem to delay the onset of aggregation (Supplementary Figure 8A). Interestingly, the aggregation curve of PSMα1 at 8.6 μM follows a typical sigmoidal curve, whereas at 77.7 μM a stepwise aggregation curve is observed (Supplementary Figure 8A, Figure 1C).

**Figure 2.**
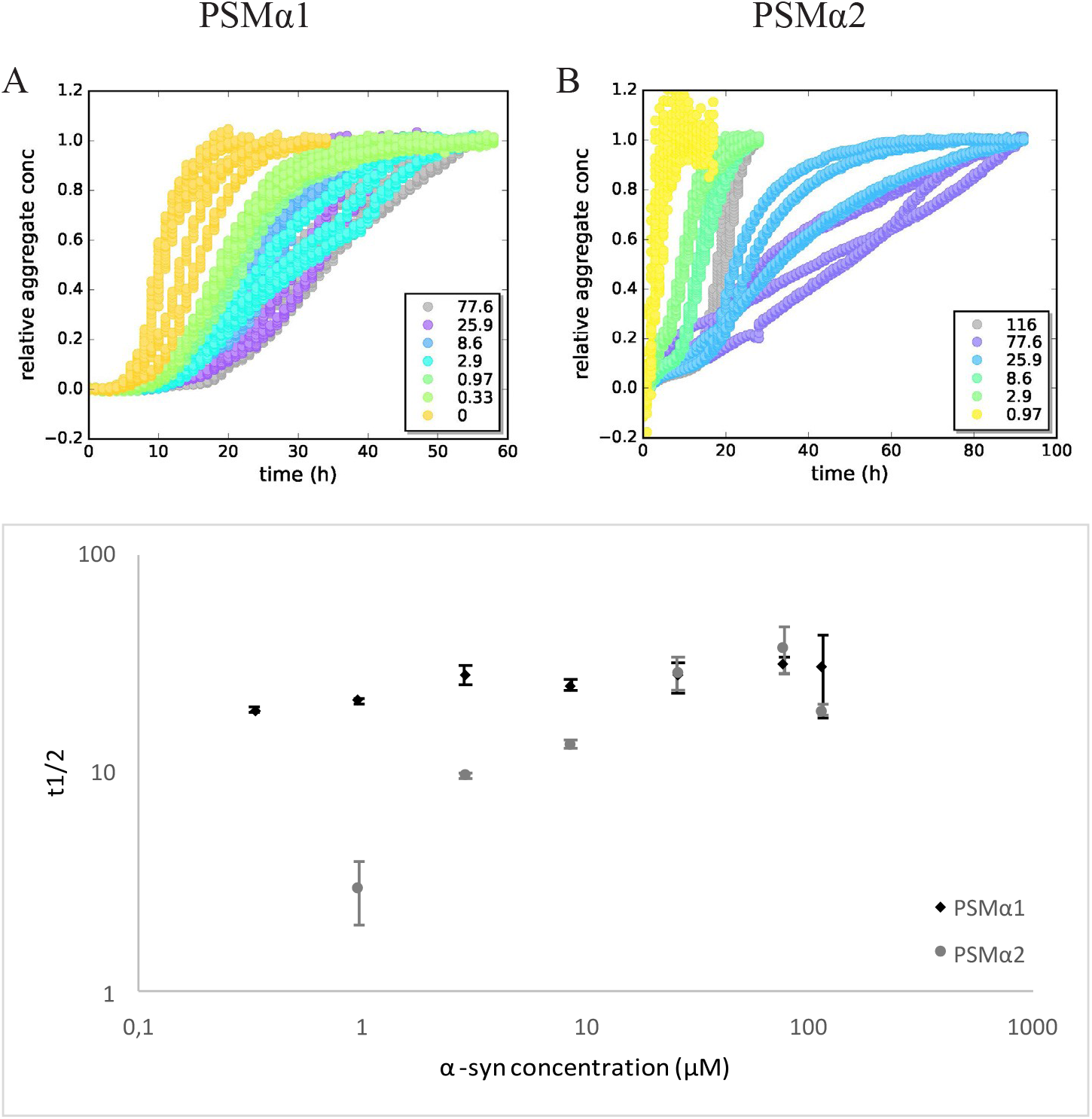
PSMα1 and PSMα2 display different α-syn affinities. The concentrations of A. PSMα1 and B. PSMα2 were kept constant at 8.6 μM and the concentration (μM) of monomeric α-syn varied in Tris buffer. The plots of t1/2 as a function of α-syn concentration in the presence of PSMα1 reveals lower α-syn monomer dependency than for PSMα2. The plots show the averages and standard deviations of replicates.

For α-syn aggregation induced by PSMα1, a low degree of monomer-concentration dependency is observed (Figure 2). This could indicate that the number catalytic sites for heterogenic primary nucleation are limiting, or that product release is the rate-limiting step (*42*).

For PSMα2 a bipahsic aggregation curve can be observed for all concentrations of α-syn, especially obvious at 25.9 and 8.6 μM. This stepwise process suggest that the process becomes physically restricted, e.g. due to restricted diffusion from gel formation or accumulation of intermediates at interfaces (Supplementary Figure 8B) (*43*). At high α-syn monomer concentrations, the aggregation is less catalyzed. Indeed, from the previous experiment, it can also be observed that PSMα2-induced catalysis of α-syn aggregation is less effective when the monomer to peptide concentration exceeds a ratio of 1:1.

The peptides are all potent inducers of α-syn aggregation, PSMα2 seemingly by heterogenous primary nucleation and PSMα1 with mechanisms yet to be elucidated. The PSMα peptides seemingly have different affinities for α-syn and likely promote oligomer species formation along the aggregation pathway.

### PSMαs do not form heteromolecular aggregates with α-syn

To gain insight into whether PSMα peptides formed heteromolecular aggregates with α-syn, we analyzed whether any co-aggregated materials had formed by Mass Spectrometry. Different PSMα-α-syn samples after aggregation were size-separated by SDS PAGE and the <10, 37, and 120 kDa bands were excised, subjected to in-gel digestion and analyzed with mass spectrometry. PSMα was detected in the samples containing the peptide alone (<10 kDa bands). All other bands showed presence of α-syn but not PSMα (Supplementary Figure 9). We thus hypothesize that the PSMαs induce α-syn aggregation by transient or weak interactions but do not become incorporated in the final aggregates.

### PSMα -peptide-induced-α-syn aggregates seed α-syn in cells

To investigate whether the formed α-syn aggregates could induce seeding in cells, the co-aggregated materials were added to HEK 293T cells expressing the A53T mutant form of α-syn fused to GFP. After 48 hours, the cells were fixed and stained for phosphorylated α-syn. All of the imaged wells showed some aggregate formation in the cells. The quantification of the GFP aggregates is less accurate than the quantification of the phosphorylated aggregates owing to the lower signal-to-noise ratio. The different PSMα-α-syn aggregates showed differences in their ability to induce phosphorylation of aggregates in cells (Figure 3, Supplementary Figure 10). This could indicate that the concentration of PSMα affects the seeding potential of the formed α-syn aggregate. When cells were treated with α-syn incubated in the absence of PSMα no phosphorylated aggregates could be seen.

**Figure 3.**
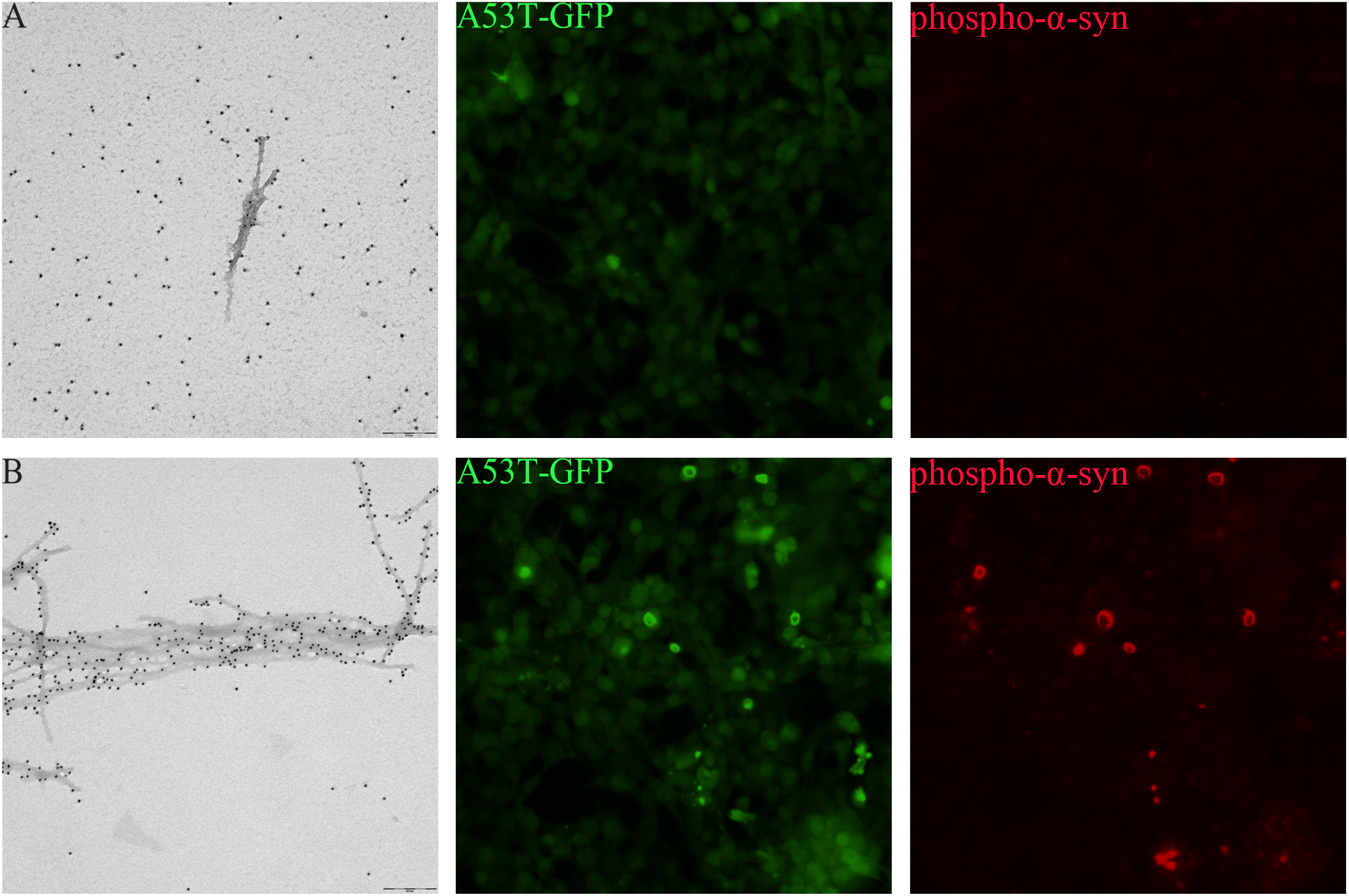
PSMα peptides induce at α-syn fibril formation which seeds α-syn in cells. TEM micrographs at 60000 magnification of 25 μM α-syn incubated with A. 1.1% DMSO or B. 8.6 μM PSMα1 peptide in 1.1% DMSO. The samples were labelled by syn211 and 10 nm immunogold. These samples were sonicated and added 1:10 to HEK cells expressing A53T α-syn fused to GFP. 48 hours post treatment, the cells were fixed and immunolabeled for phosphorylated α-syn. Scale bar = 200 nm

## Discussion

α-Syn is thought to be an intrinsically disordered monomer *in vivo,* which forms aggregates under disease conditions. After the formation of the initial aggregates, rapid amplification of the aggregate mass can occur by autocatalytic secondary nucleation or by monomer addition to existing aggregates in an elongation process (*44*). What triggers the formation of the initial seeds, however, remains unclear. The energy barrier for *de novo* seed formation is high and homogenous primary nucleation, whereby multimeric species arise from monomer species in solution. At neutral pH, under quiescent conditions, primary nucleation is undetectable for α-syn, at least over typical experimental time-frame of a few weeks (*45*). Heterogenous primary nucleation, on the other hand, occurs on the surface of other substances such as polystyrene plates or nanoparticles (*46*) or lipid vesicles (*40*), or at the air-liquid interface (*47*). *In vivo*, primary nucleation is thought to be mainly heterogenous (*44*). Extrinsic inducers of aggregation would present an explanation for the role of the environment in idiopathic PD. In our current work, we identified the bacterial amyloids, formylated PSMα peptides produced by *S. aureus,* as potent catalyzers of α-syn aggregation.

The α-Syn aggregation mechanism is highly dependent on solution conditions such as pH and ionic strength, owing to its heterogenous charge distribution (*45*). At a mildly acidic pH, the overall net negative charge of α-syn is reduced compared to at neutral pH, leading to a decrease in electrostatic repulsion between the positively charged N-terminal and negatively charged C-terminal of different α-syn molecules and an increase in aggregation propensity. High salt concentrations, on the other hand, have been shown to screen electrostatic attractions between the termini within or between monomers in solution and on surfaces, resulting in a retardation of aggregation (*48*). In this study we show that the PSMα peptides rapidly induce α-syn aggregation even at neutral pH and moderate ionic strength overcoming a high energy barrier (*44*). Even though all of the examined PSMα peptides accelerate α-syn aggregation, the mechanisms by which they do so seem to differ. PSMα2 seems to catalyze α-syn aggregation through primary heterogeneous nucleation with the rate of aggregation being dependent on the ratio of PSMα2 peptide to α-syn. In contrast, a model of only primary nucleation and elongation cannot explain the kinetic data observed for PSMα1 induced α-syn aggregation. PSMα1 differs from PSMα2 by only 3 amino acids and seems to have a different effect on α-syn aggregation as indicated by the low monomer-concentration dependency of aggregation. The PSMα peptides themselves have been shown to have different pH and concentration dependent aggregation kinetics (*41*). Further work may elucidate the molecular origin of these differences.

Peripheral α-syn expression is well documented and several studies have shown α-syn propagation and seeding from the periphery to the CNS (*3–6*). Thus, we studied whether the PSMα-induced α-syn aggregates could induce seeding in cells. Indeed, upon direct addition to a cell line, the formed α-syn aggregates also catalyzed α-syn aggregation in the cells and increased phosphorylation of the aggregates. Again, we could see differences in this catalytic effect depending on the presence of the different PSMα peptides and their concentration. Likewise in the aggregation kinetics, we see a clear dependence on the ratio of peptide to α-syn.

Our current work identifies PSMα peptides as potent extrinsic inducers of α-syn aggregation and potential triggers of α-syn pathology. Our current work also highlights the concentration-dependent interactions of the PSMα peptides and α-syn monomer on the aggregation kinetics and the triggering of aggregation in cells. This supports that underlying genetic predispositions and bacterial load could affect the pathological outcome. Previous studies have shown propagation of aggregated α-syn from peripheral tissues to the CNS. Our work identifies a mechanism by which the original seeds of α-syn could arise in the periphery. Figure 4 illustrates how a *S. aureus* infection, could trigger a cascade of NETosis and increased PSMα production catalysing α-syn aggregation in peripheral tissues. With *S. aureus* infections continuing to rise globally, with the elderly population especially affected, host-pathogen interactions will gain further interest from the research and medical fields. The current results could have significant implications for the understanding of *S. aureus* in initiation of α-syn pathology in different synucleinopathies, such as PD.

**Figure 4.**
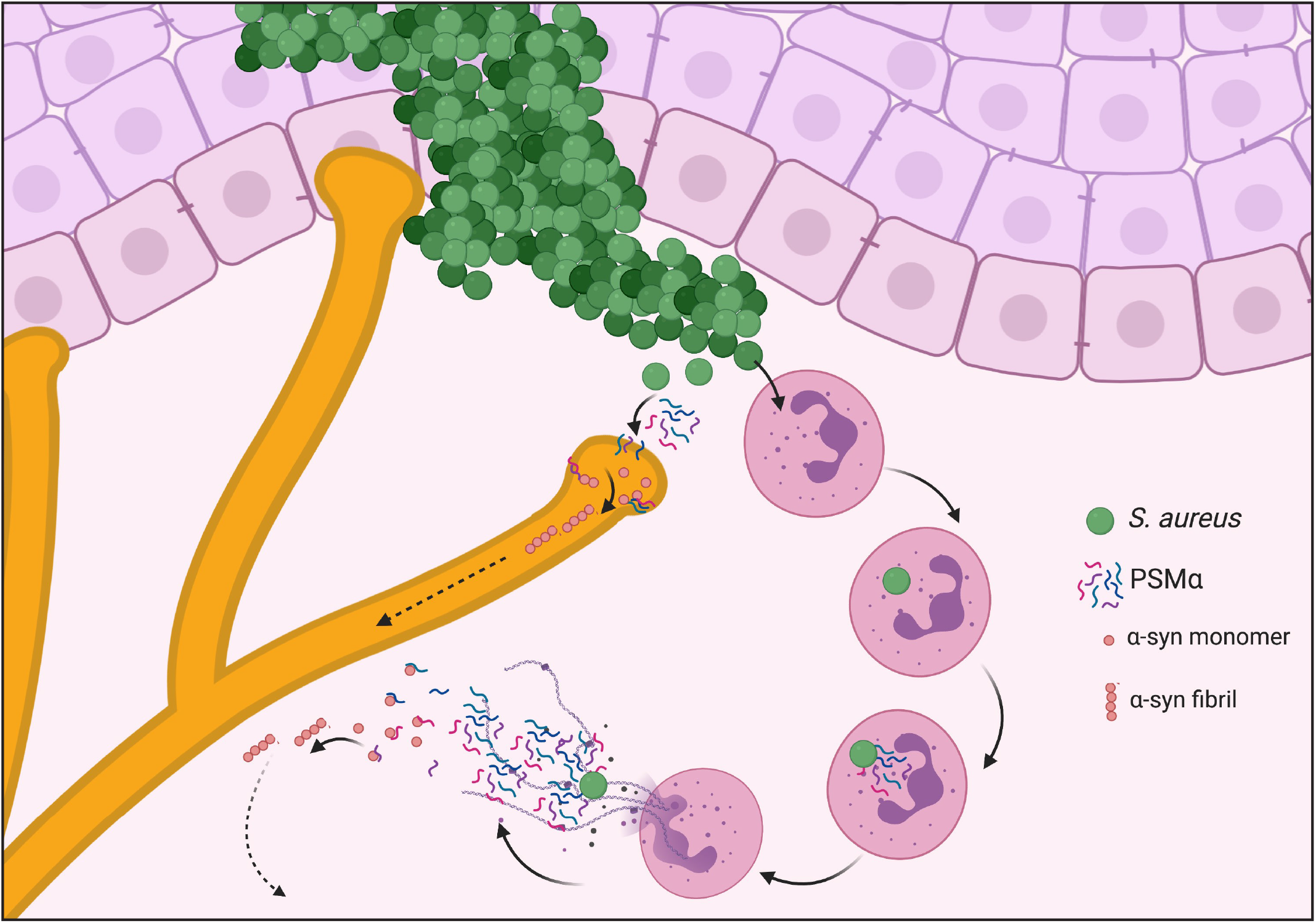
Schematic illustration of *S. aureus* resulting in PSMα-induced α-syn aggregation. Invading *S. aureus* produce PSMα peptides chemoattracting neutrophils. Neutrophils phagocytose S. aureus bacteria, whereby PSMα expression is upregulated leading to phagosome escape and NETosis (21, 22, 34). The high concentrations of PSMα peptides interact with extracellular α-syn causing aggregation by primary nucleation leading to the formation of fibrils which can propagate and seed α-syn in other cells (this study). PSMα peptides also form pores in sensory neurons. The PSMα peptides can then interact with intracellular α-syn.

## Acknowledgements

The authors would like to thank Johan Bylund for his generous contribution of an aliquot each of PSMα2 and PSMα3 for the pilot experiments. The authors would also like to thank LBIC and Lina Gefors for guidance with the Transmission Electron Micrsocopy, Birgitta Frohm for guidance with the Size Exclusion Chromatography and EMC microcollections for production of the PSMα peptides. BioRender was used for the illustration in Figure 4.

## Author Contributions

CH, JYL and SL conceived of and planned the project. KB purified α-syn and performed the in-gel digestion of size-separated materials and subsequent mass spectrometry analysis. LO performed the SDS gel separation of materials. AS produced the HEK A53T GFP cell line. Remaining experiments were performed by CH with continuous input from SL and JYL. All authors have contributed to and edited the manuscript text and figures.

