## Supplementary Figures for "The bacterial amyloids phenol soluble modulins from *Staphylococcus aureus* catalyze alpha-synuclein aggregation"

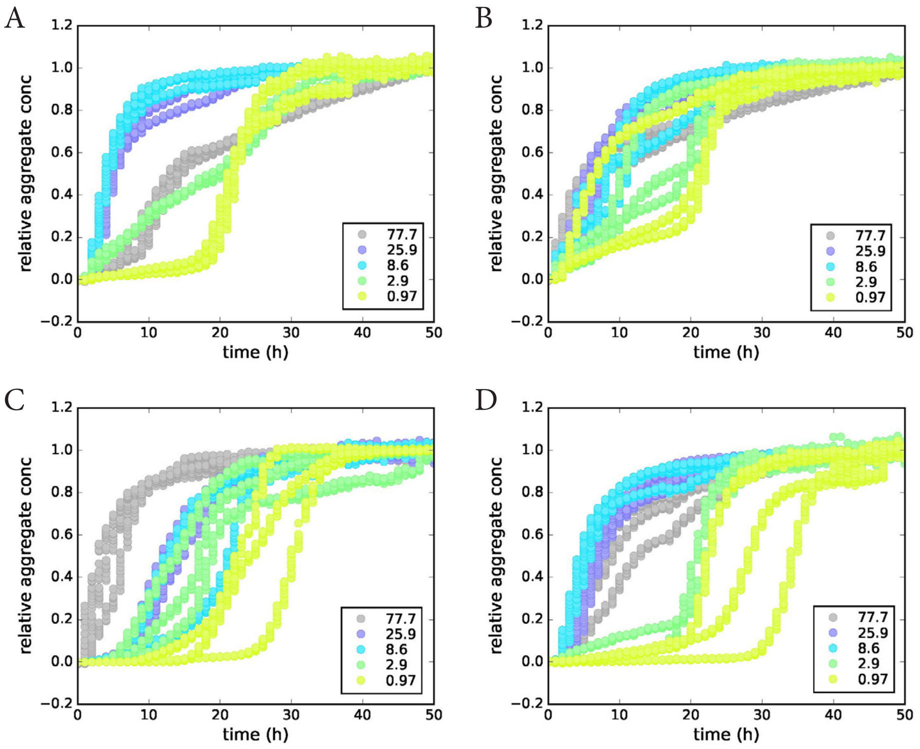


Supplementary Figure 1. Aggregation kinetics in MES buffer. The normalized ThT fluorescence intenisities of α-syn aggregated in the presence of varying concentrations (μM) of A. PSMα1 B. PSMα2 C. PSMα3 and D. PSMα4 as a function of time.

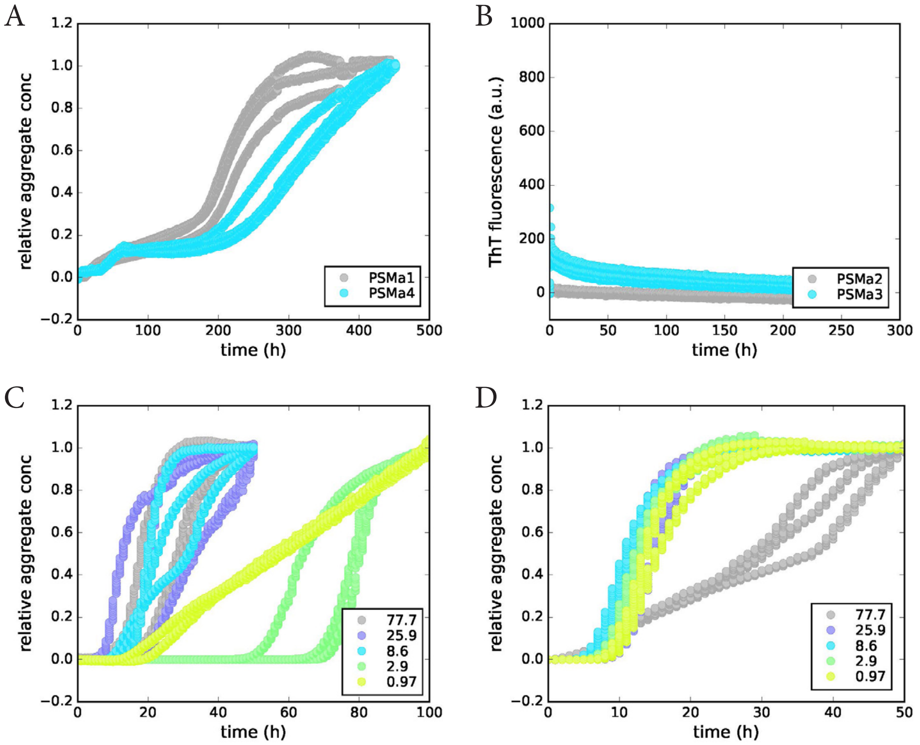


Supplementary Figure 2. PSMα peptides induce rapid aggregtion of α-syn. A. The normalized ThT fluorescence intensity of pure PSMα1 and PSMα4 as a function of time. B. The non-normalized ThT fluorescence intensity of PSMα2 and PSMα3 as a function of time. The normalized ThT fluorescence intenisities of 25 μM monomeric α-syn aggregated in the presence of varying concentrations (μM) of C. PSMα3 and D. PSMα4 as a function of time. Even low concentrations of any PSMα accelerate α-syn aggregation. Pure monomeric α-syn did not aggregate under these conditions.

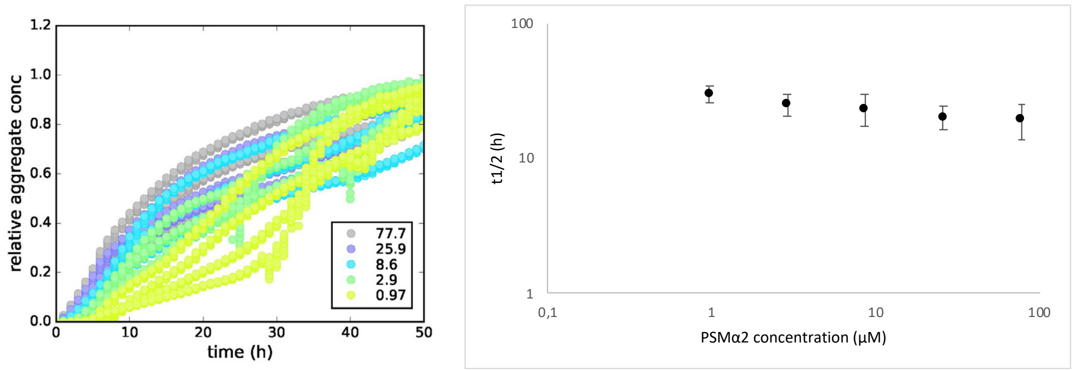


Supplementary Figure 3. DMSO does not affect concentration dependenace of PSMα2-induced α-syn aggregation. Each dilution of PSMα2 was supplemented with additional DMSO to correspond to the concentration of DMSO (8.9% v/v) in the most conentrated peptide sample. The different dilutions were added to α-syn in Tris buffer and ThT fluorescence measured. The t1/2 was plotted against the concentration of PSMα2.

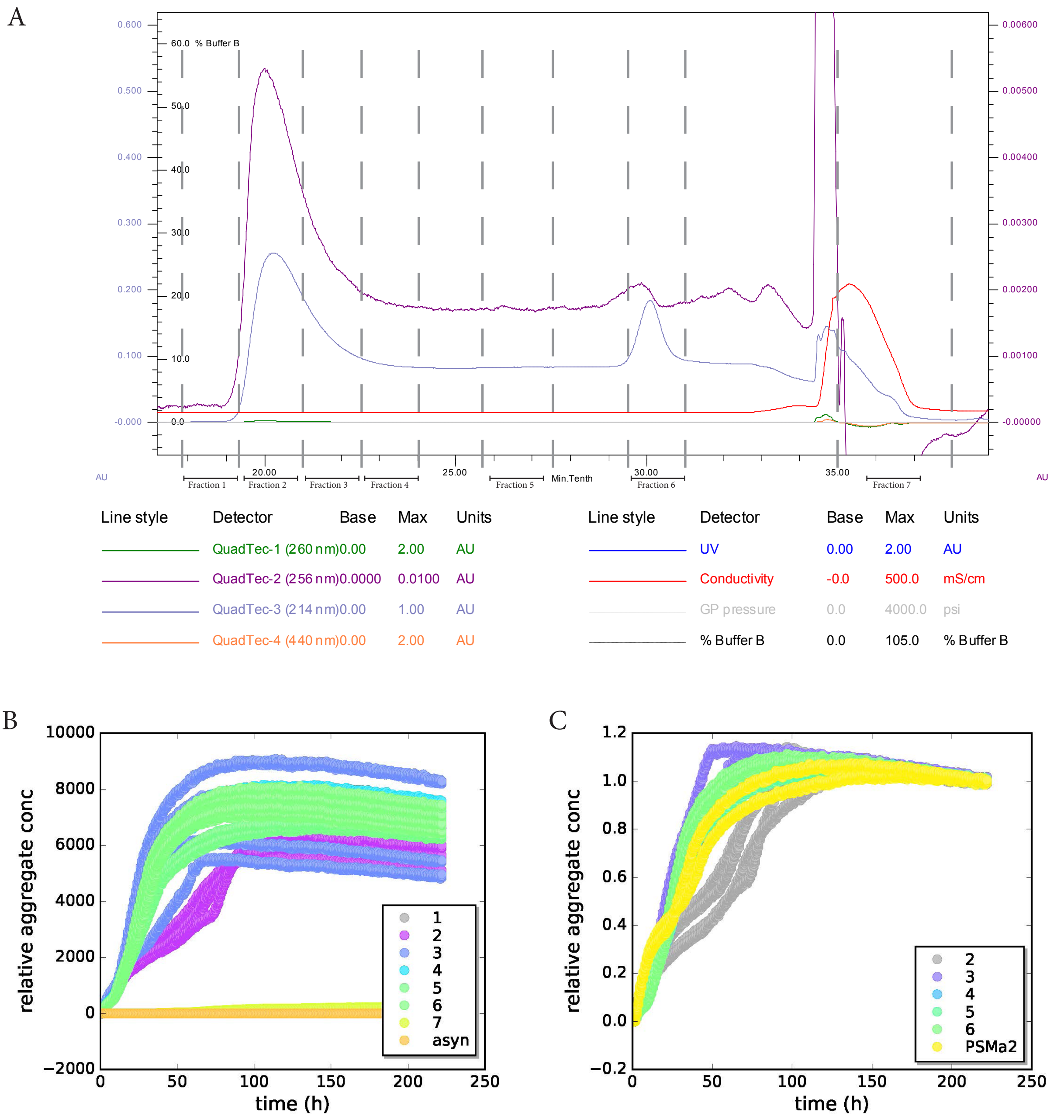


Supplementary Figure 4. All peptide-containg fractions of purified PSMα2 induce rapid α-syn aggregation. PSMα2 was disolved in 6M GuHCl and purified using size exclusion chromatography. A. The absorbance at 214 and 256 nm was monitored and several fractions collected. The collected samples were co-incubated with 25 µM freshly purified α-syn, 2 µM ThT and 0.01% NaN3. B. The non-normalized kinetics reveal that all peptide containg fractions (2-6) induce α-syn aggregation. α-syn incubated with fractions not containg peptide (1 and 7) as well as pure α-syn did not aggregate during the course of this experiment. A sample of PSMα2 dissolved in DMSO and not re-purified was used as a control. C. The normalized ThT plot reveals similar kinetics for the purified fractions of PSMα2 and the whole peptide sample.

Supplementary Figure 5. PSMα induce asyn fibril formation. TEM micrographs at 60000 magnification of asyn or PSMα peptides aggregated on their own or together. The samples were labelled by syn211 and 10 nm immunogold. A. PSMα1 only B. α-syn C. PSMα2 only D. α-syn and PSMα2. E. PSMα3 only F. α-syn and PSMα3. G. PSMα4 only H. α-syn and PSMα4
Scale bar = 200 nm

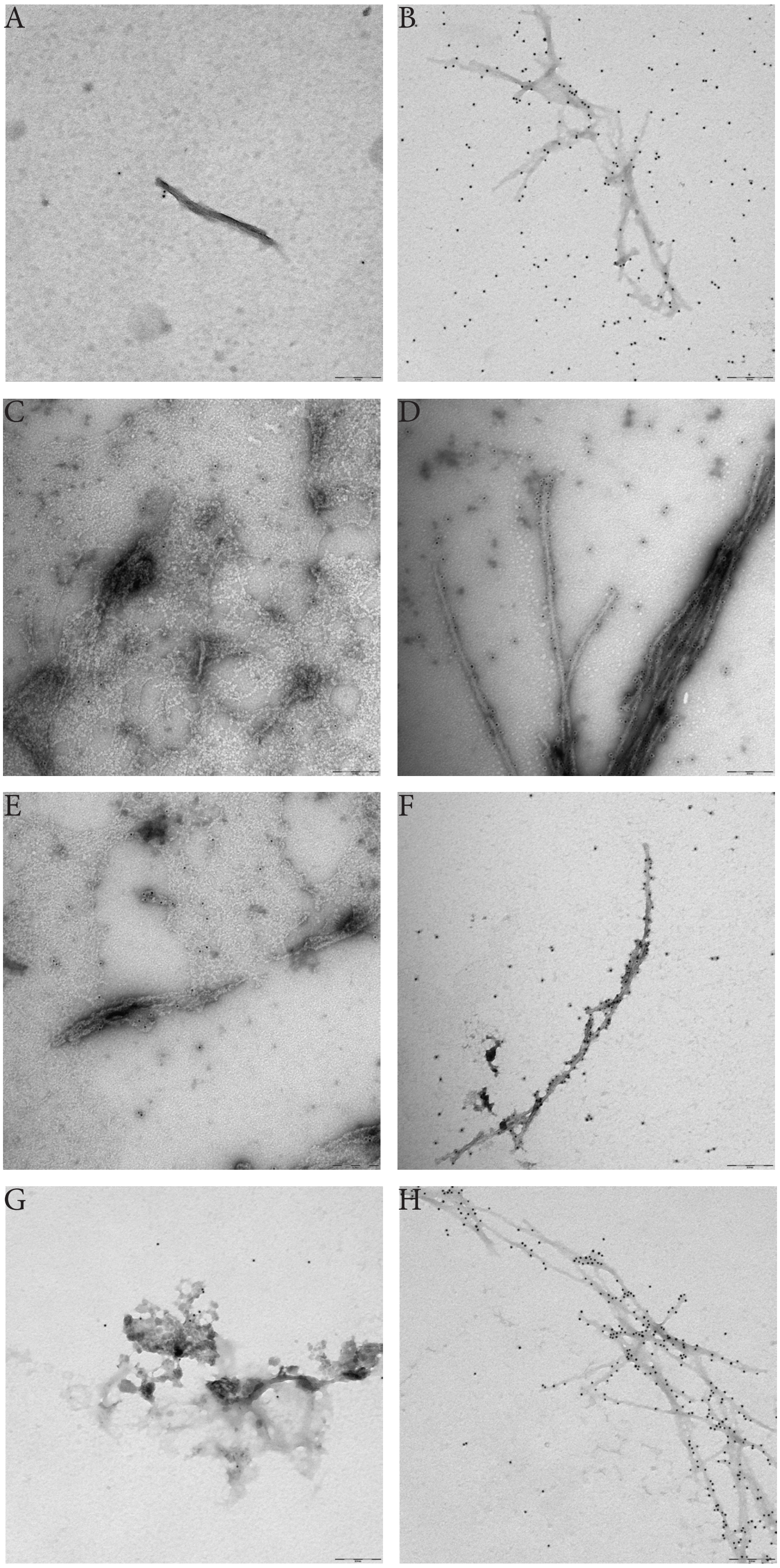

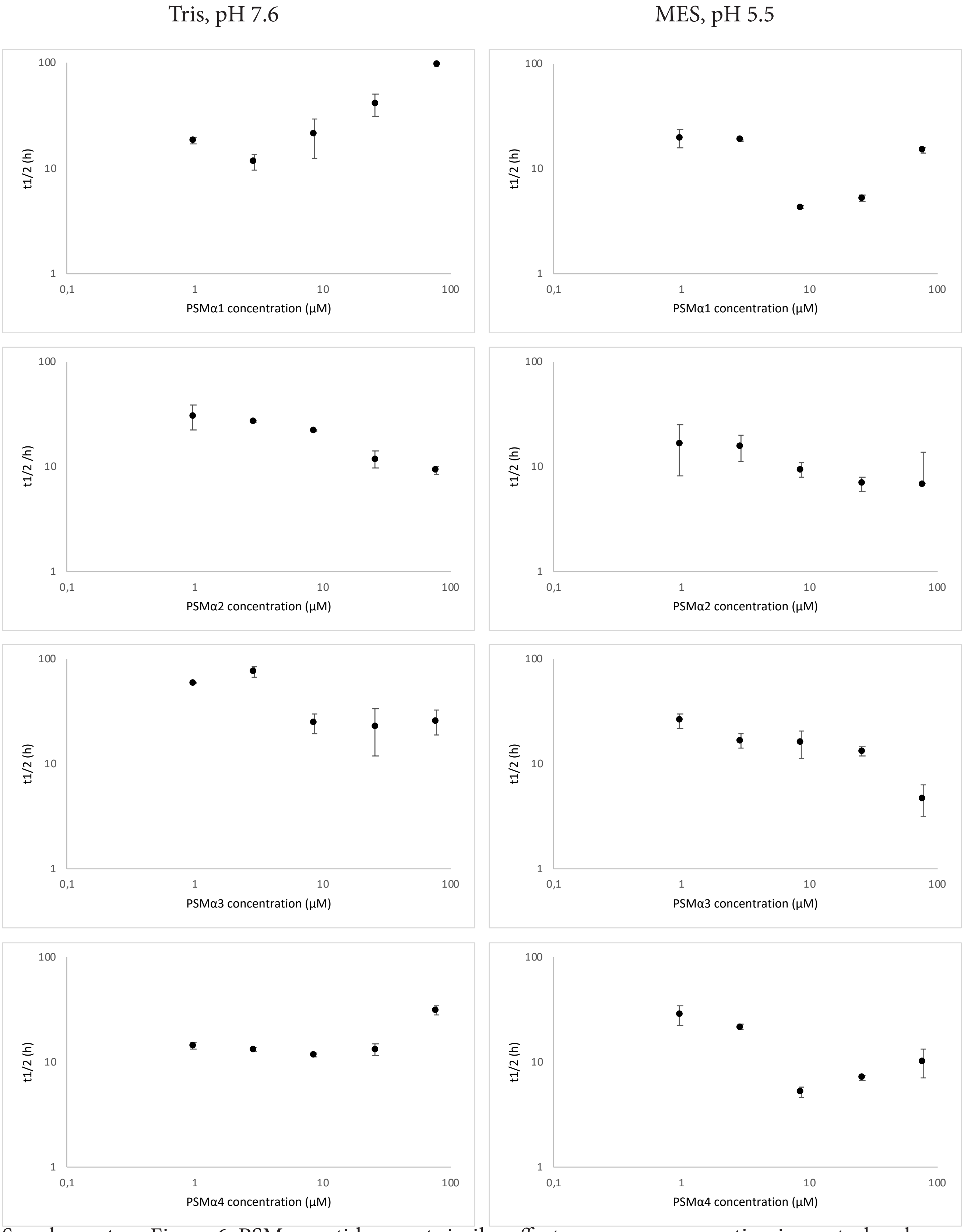


Supplementary Figure 6. PSMα peptides exert similar effects on α-syn aggregation in neutral and slightly acidic buffers. The t1/2 of α-syn aggregated in the presence of different concentrations of PSMα peptides were extracted and plotted against the concentrations of the peptides. The averages and standard deviations from replicates from a single experiment are shown.

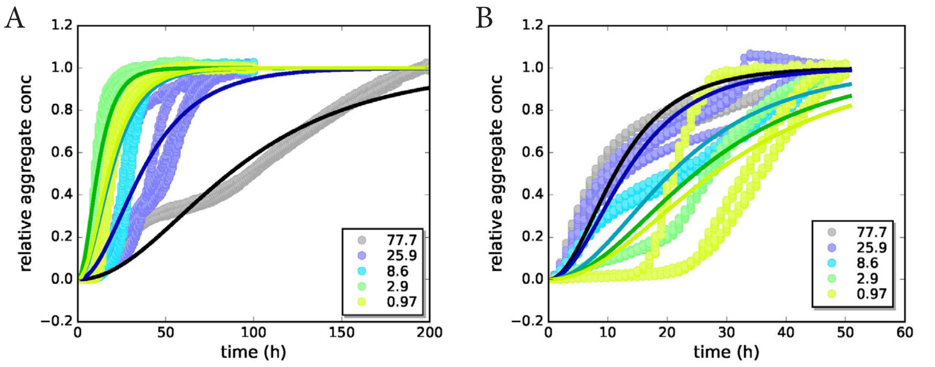


Supplementary Figure 7. The kinetic curves of α-syn aggregation induced by varying concentrations of A. PSMα1 and B. PSMα2 were fitted to a model of nucleation elongation. The model fits the data of PSMα2-induced aggregation well, indicating PSMα2 likely catalyzes α-syn aggregation by heterogeneous primary nucleation. However, the model does not fit the data of PSMα1-induced aggregation, indicating there could be other mechanisms at play.

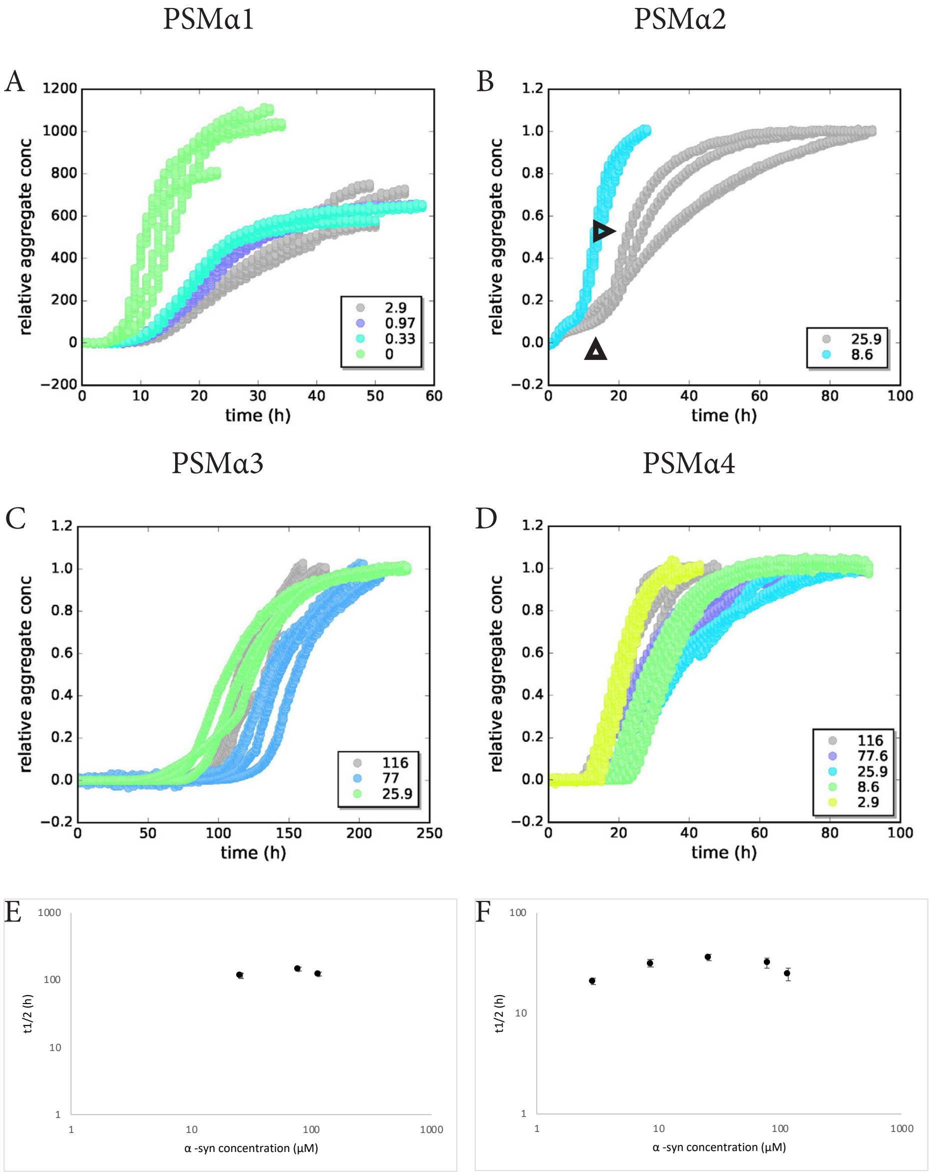


Supplementary Figure 8. Aggregation kinetics of α-syn in the presence of PSMα peptides. The ThT fluorescence was monitored when concentrations of A. PSMα1 B. PSMα2 C. PSMα3 and D. PSMα4 were kept constant at 8.6 µM and the concentration (µM) of monomeric α-syn varied in Tris buffer. The non-normalized ThT intensities indicate pure PSMα1 aggregates quicker than in the presence of low concentrations of α-syn. α-syn aggregated in the presence of PSMα2 shows multistep aggregation curves indicative of oligomer formation (arrowheads). PSMα3 did not induce aggregation of lower concentrations of α-syn. The t1/2 was plotted as a function of α-syn concentration for E. PSMα3 and F. PSMα4

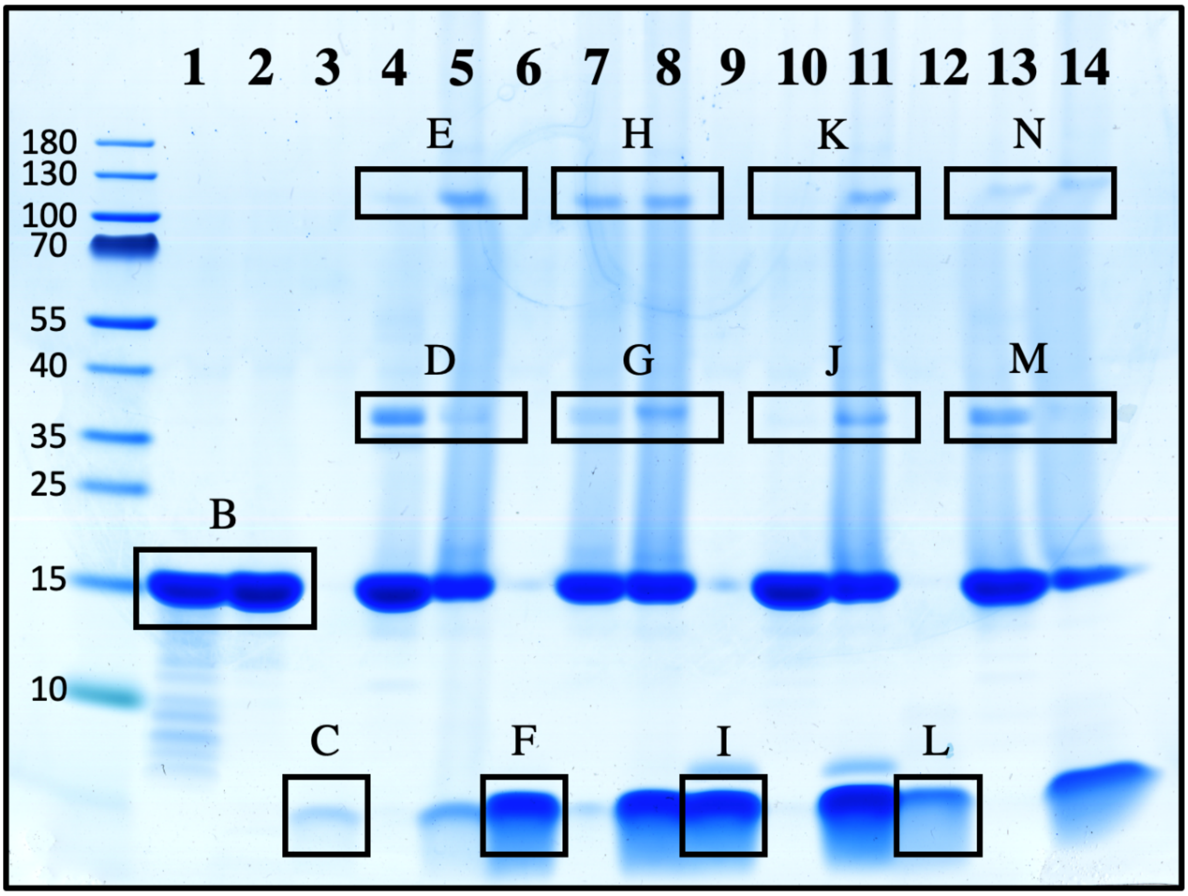


| Gel Lane | Sample | Gel band/Spectrum |
|---|---|---|
| 1 | Standard | |
| 2 | α-syn+ 0.9% DMSO | B |
| 3 | α-syn + 8.9% DMSO | |
| 4 | 77.7 µM PSMα1 | C |
| 5 | 25 µM α-syn + 0.97 µM PSMα1 | D and E |
| 6 | 25 µM α-syn + 77.7 µMPSMα1 | D and E |
| 7 | 77.7 µM PSMα2 | F |
| 8 | 25 µM α-syn + 0.97 µM PSMα2 | G and H |
| 9 | 25 µM α-syn + 77.7 µM PSMα2 | G and H |
| 10 | 77.7 µM PSMα3 | I |
| 11 | 25 µM α-syn + 0.97 µM PSMα3 | J and K |
| 12 | 25 µM α-syn + 77.7 µM PSMα3 | J and K |
| 13 | 77.7 µM PSMα4 | L |
| 14 | 25 µM α-syn + 0.97 µM PSMα4 | M and N |
| 15 | 25 µM α-syn + 77.7 µM PSMα4 | M and N |

Gel band/ Spectrum B

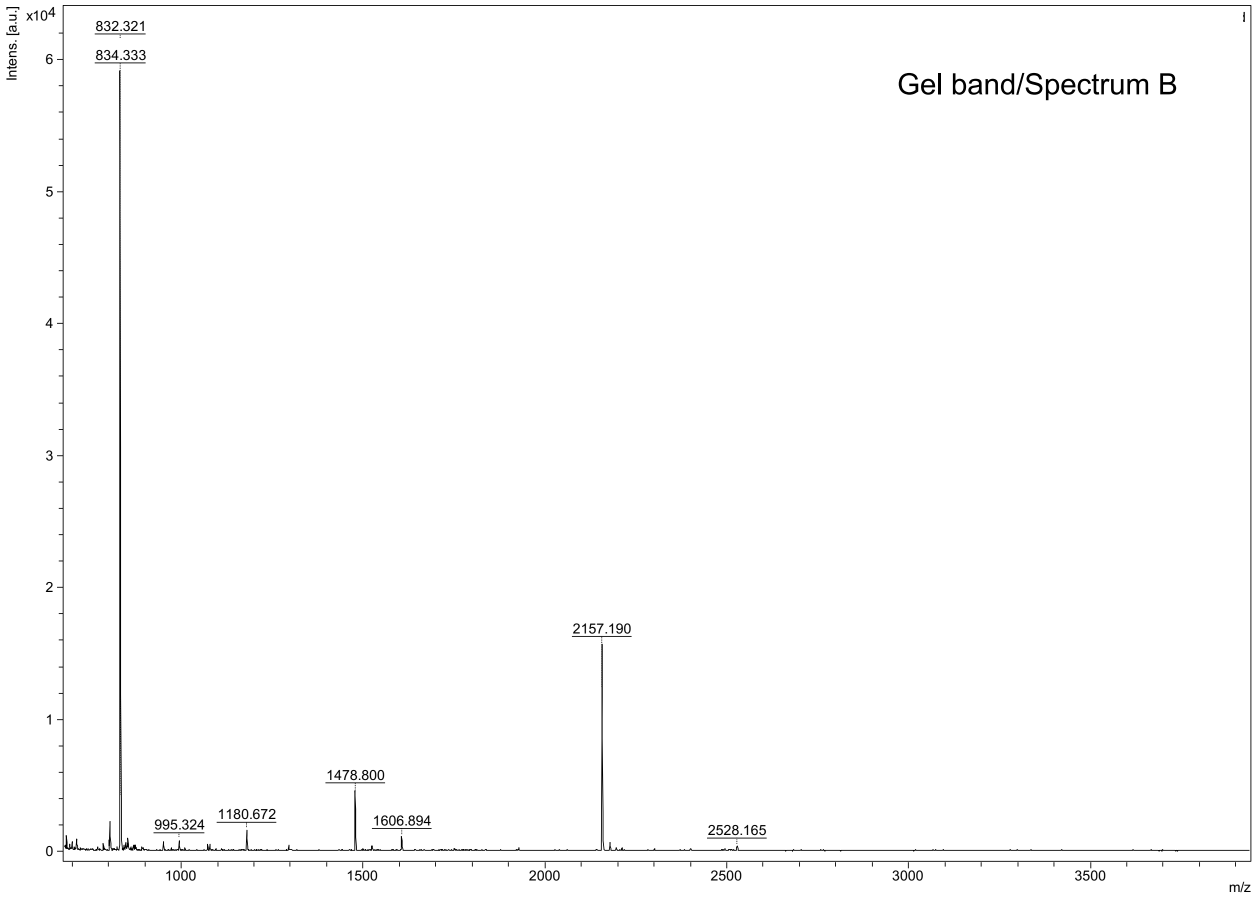

Gel band/ Spectra C, D and E (PSMα1)

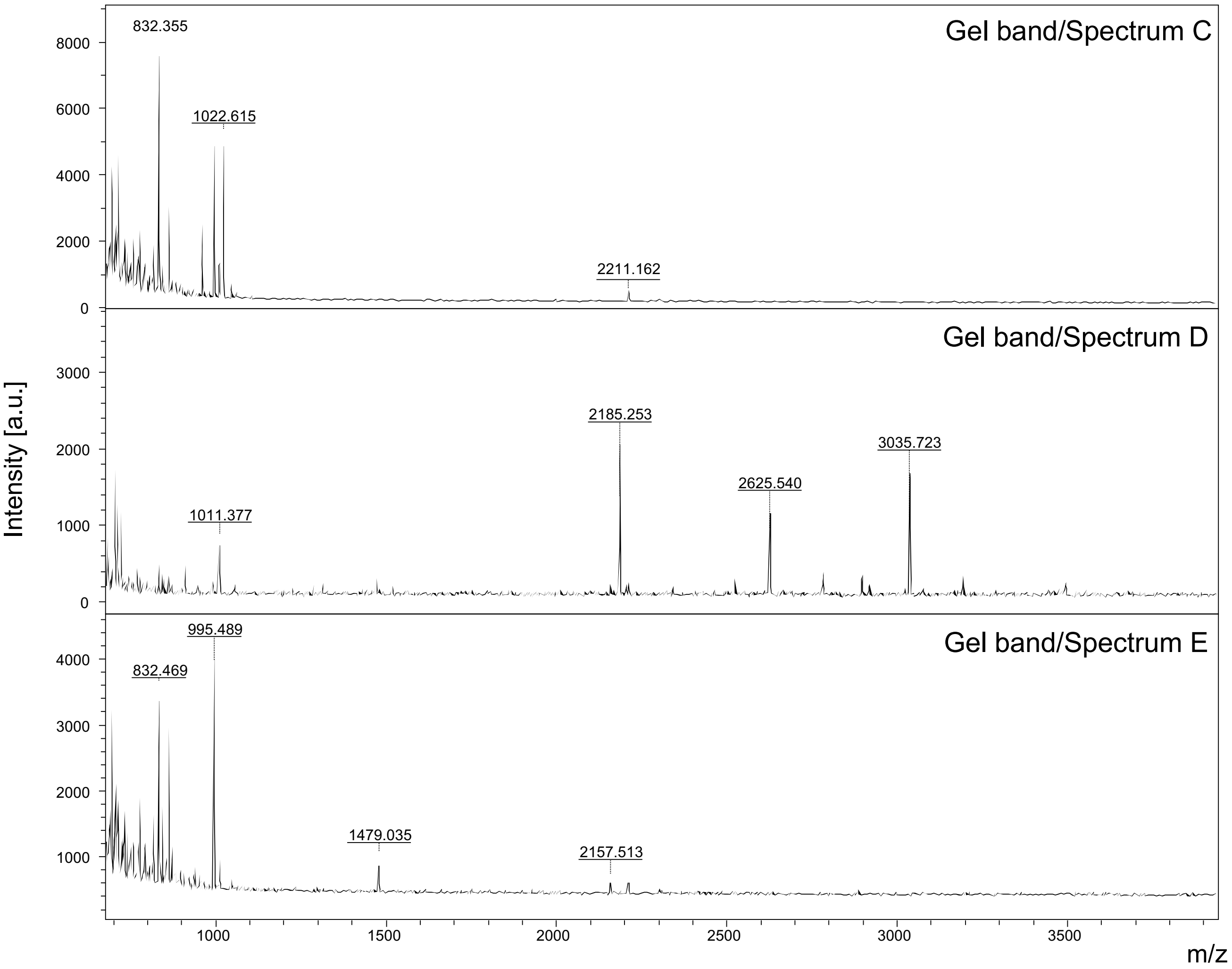


*MS spectra of in-gel digested PSMα1 samples*. **Gel band/Spectrum: C** (2 kDa band), sample with 77.7 μM PSMα1; **D** (37 kDa band) and **E** (120 kDa band) sample with pooled digests of 25 μM α-syn + 0.97 μM PSMα1 and 25 μM α-syn + 77.7 μM PSMα1. The PSMα1 peptide SLIEQFTGK (aa from 13 to 21) with a theoretical mass of 1022.55 Da was detected in gel band C (*see zoom on peptide in figure below)*

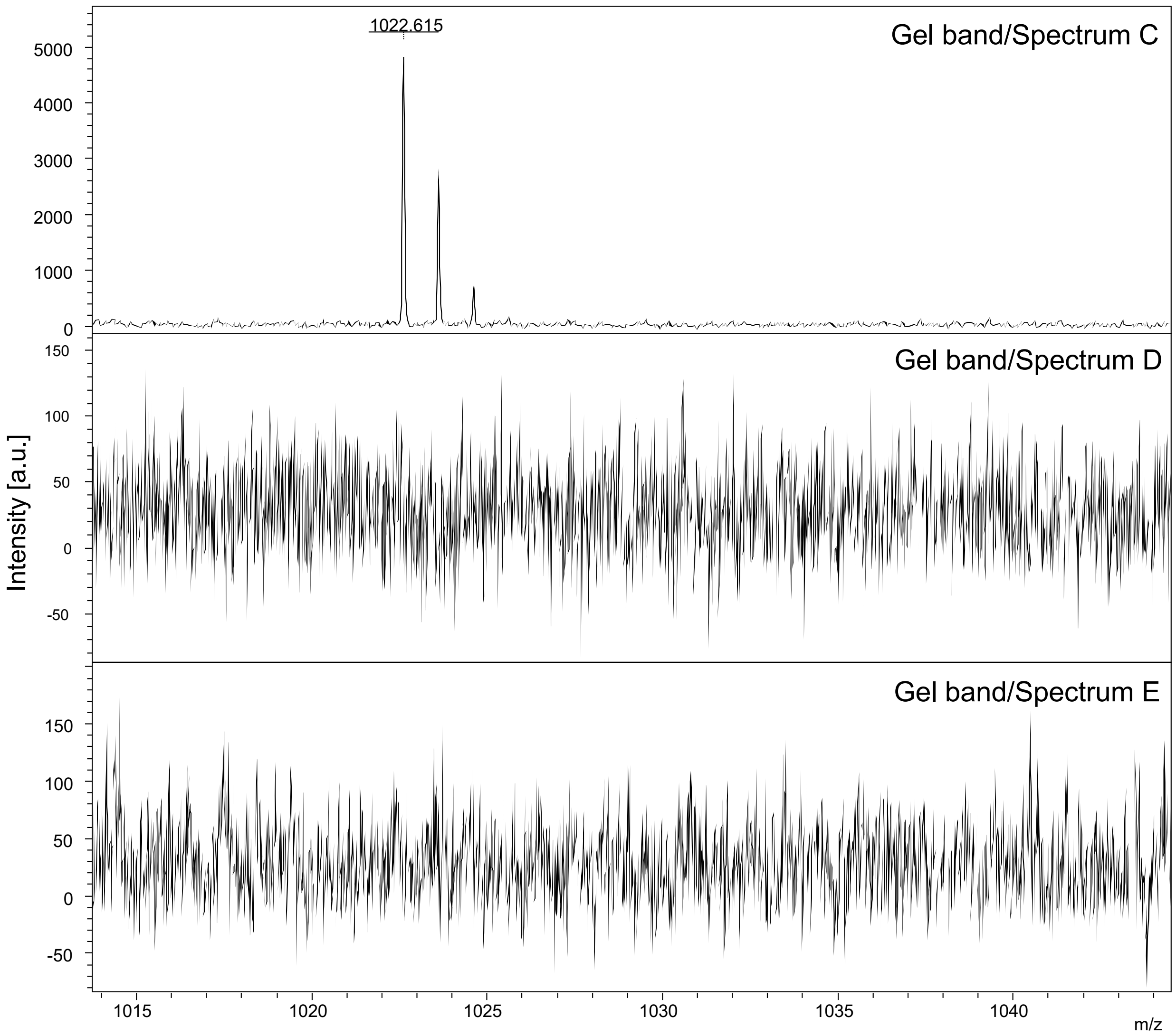

1022.615
Gel band/Spectrum C
Gel band/Spectrum D
Gel band/Spectrum E
Intensity [a.u.]
m/z

Gel band/ Spectra F, G and H (PSMα2)

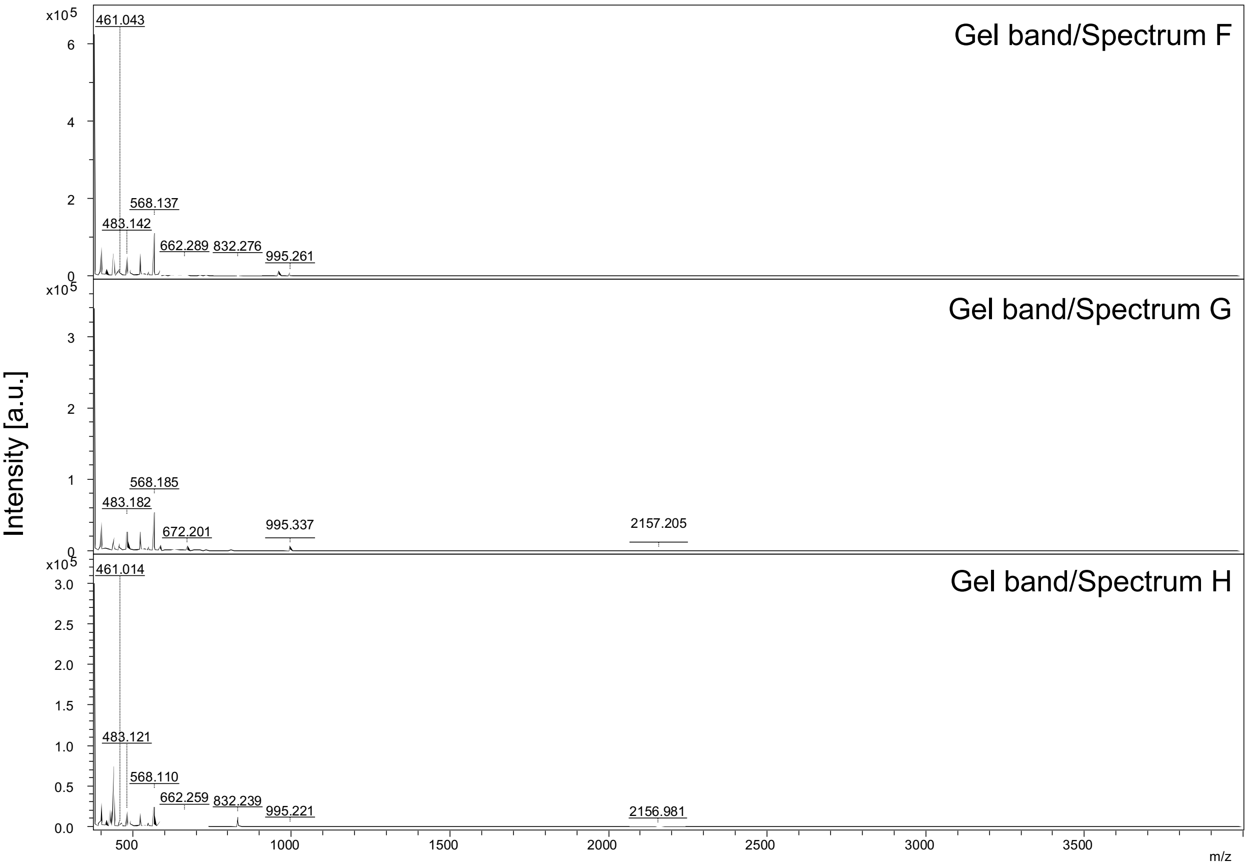


*MS spectra of in gel digested PSMα2 samples*. **Gel band/Spectrum: F** (2 kDa band), sample with 77.7 µM PSMα2; **G** (37 kDa band) and **H** (120 kDa band) sample with pooled digests of 25 µM α-syn + 0.97 µM PSMα2 and 25 µM α-syn + 77.7 µM PSMα2. The following PSMα2 peptides were detected and are presented in the zoomed spectra below: FIK (aa from 10 to 12) with a theoretical mass of 407.27 Da, FTGK (aa from 18 to 21) with a theoretical mass of 452.25 Da and GLIEK (aa from 13 to 17) with a theoretical mass of 559.34.27 Da. These were all detected in gel band F.

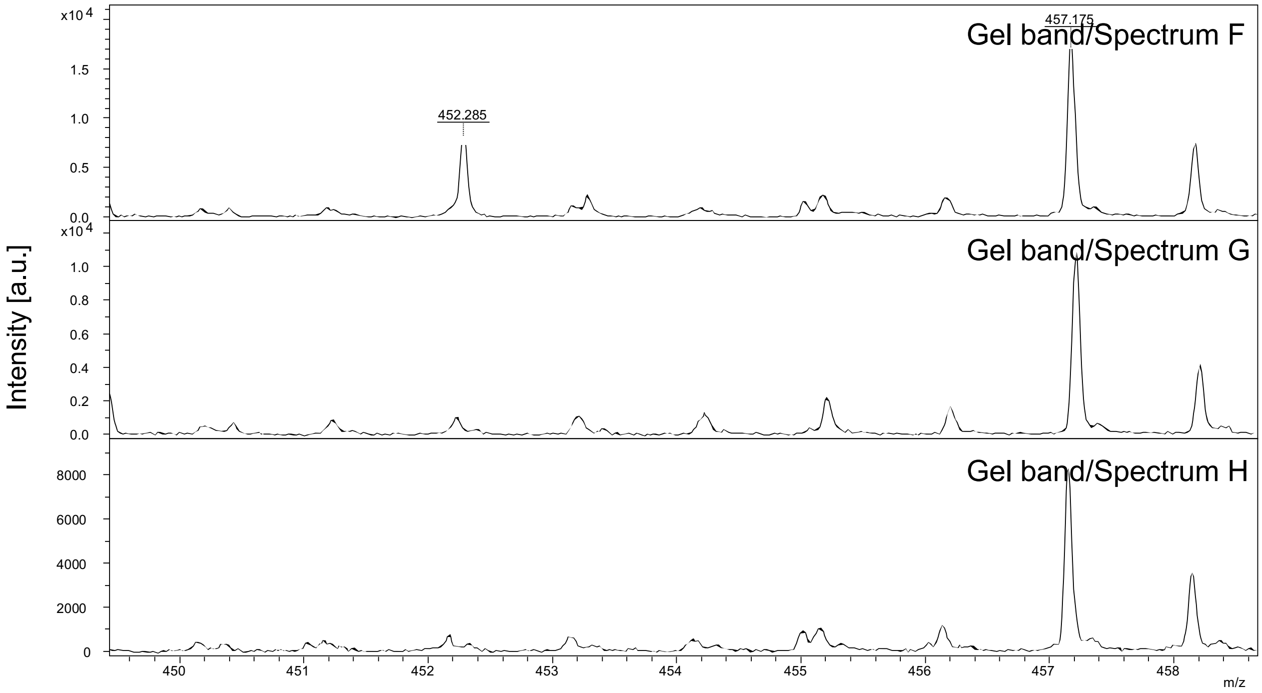
Gel band/Spectrum F
Gel band/Spectrum G
Gel band/Spectrum H
457.175
452.285
x10 4
1.5
1.0
0.5
0.0
x10 4
1.0
0.8
0.6
0.4
0.2
0.0
8000
6000
4000
2000
0
Intensity [a.u.]
450
451
452
453
454
455
456
457
458
m/z

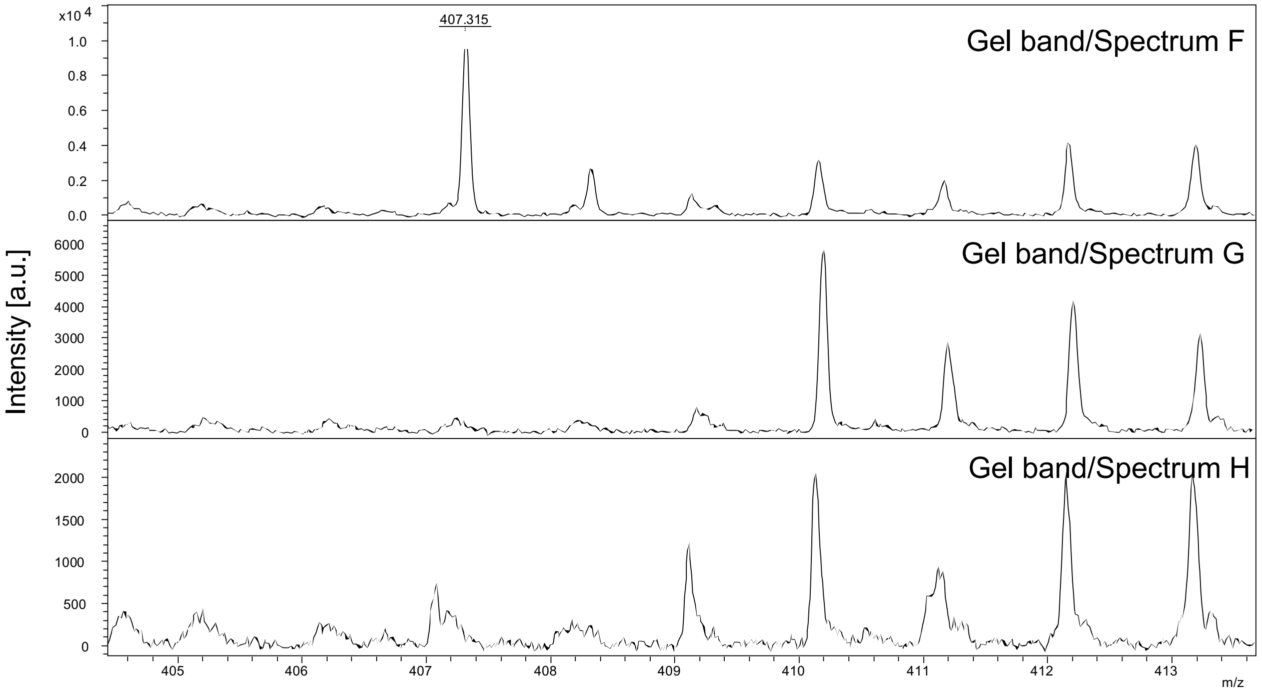
Gel band/Spectrum F
Gel band/Spectrum G
Gel band/Spectrum H
407.315
x10 4
1.0
0.8
0.6
0.4
0.2
0.0
6000
5000
4000
3000
2000
1000
0
2000
1500
1000
500
0
Intensity [a.u.]
405
406
407
408
409
410
411
412
413
m/z

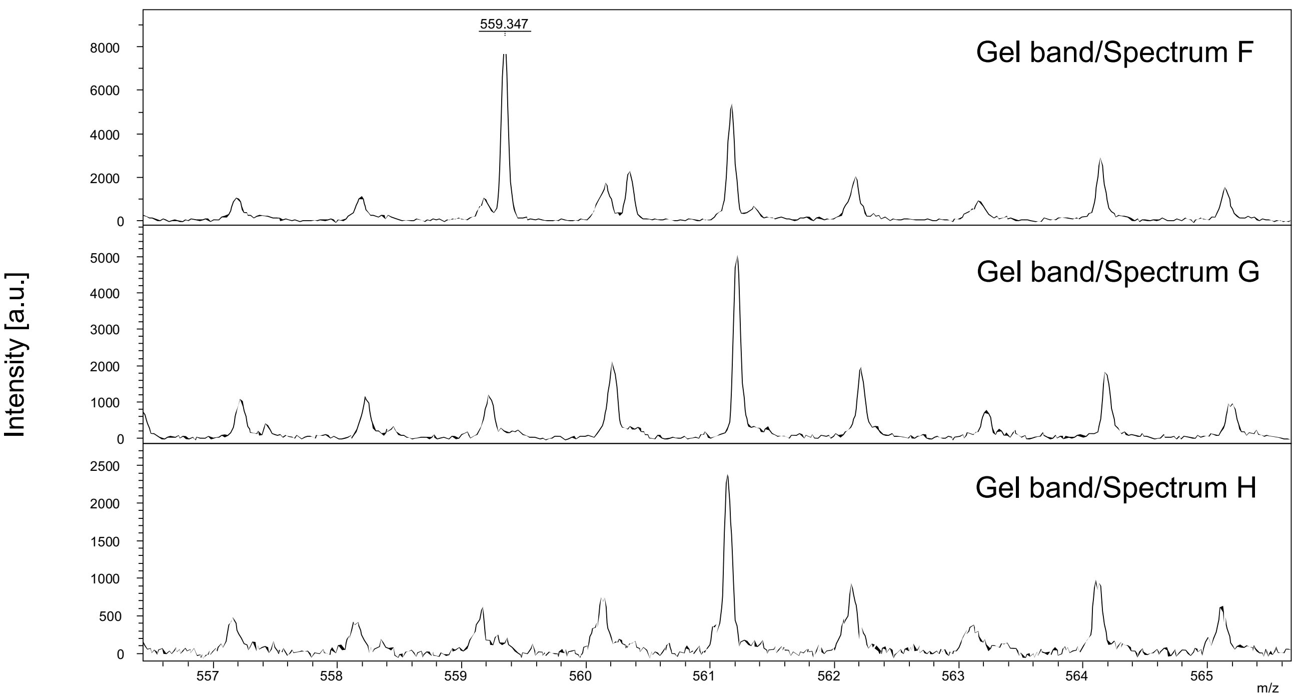

Intensity [a.u.]
559.347
Gel band/Spectrum F
Gel band/Spectrum G
Gel band/Spectrum H
m/z

Gel band/ Spectra I, J and K (PSMα3)

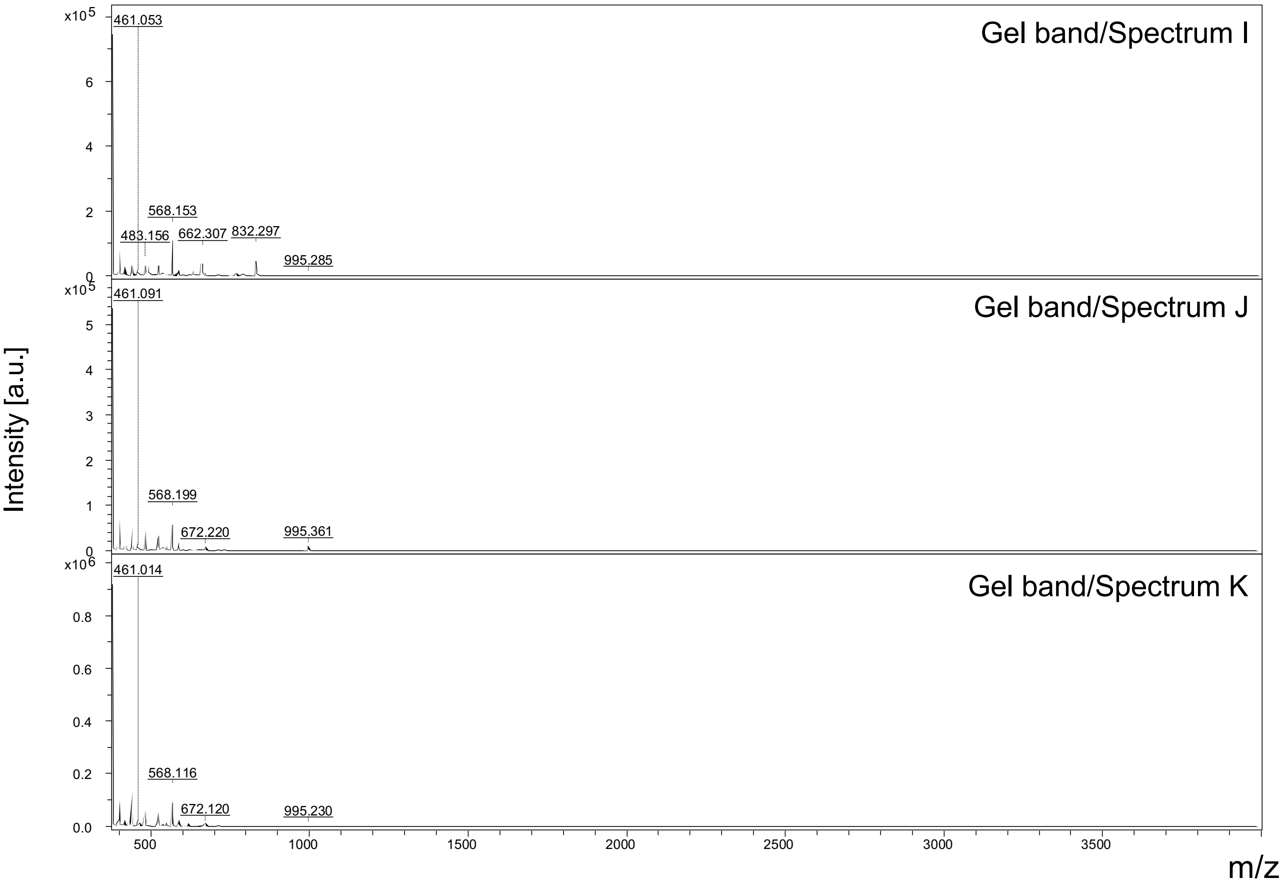


*MS spectra of in-gel digested PSMα3 samples*. **Gel band/Spectrum: I** (2 kDa band), sample with 77.7 µM PSMα3; **J** (37 kDa band) and **K** (230 kDa band) sample with pooled digests of 25 µM α-syn + 0.97 µM PSMα3 and 25 µM α-syn + 77.7 µM PSMα3. The following PSMα3 peptides were detected and are presented in the zoomed spectra below: LFK (aa from 7 to 9) with a theoretical mass of 407.27 Da, MEFVAK (aa from 1 to 6) with a theoretical mass of 752.36 Da. Both are detected in gel band I.

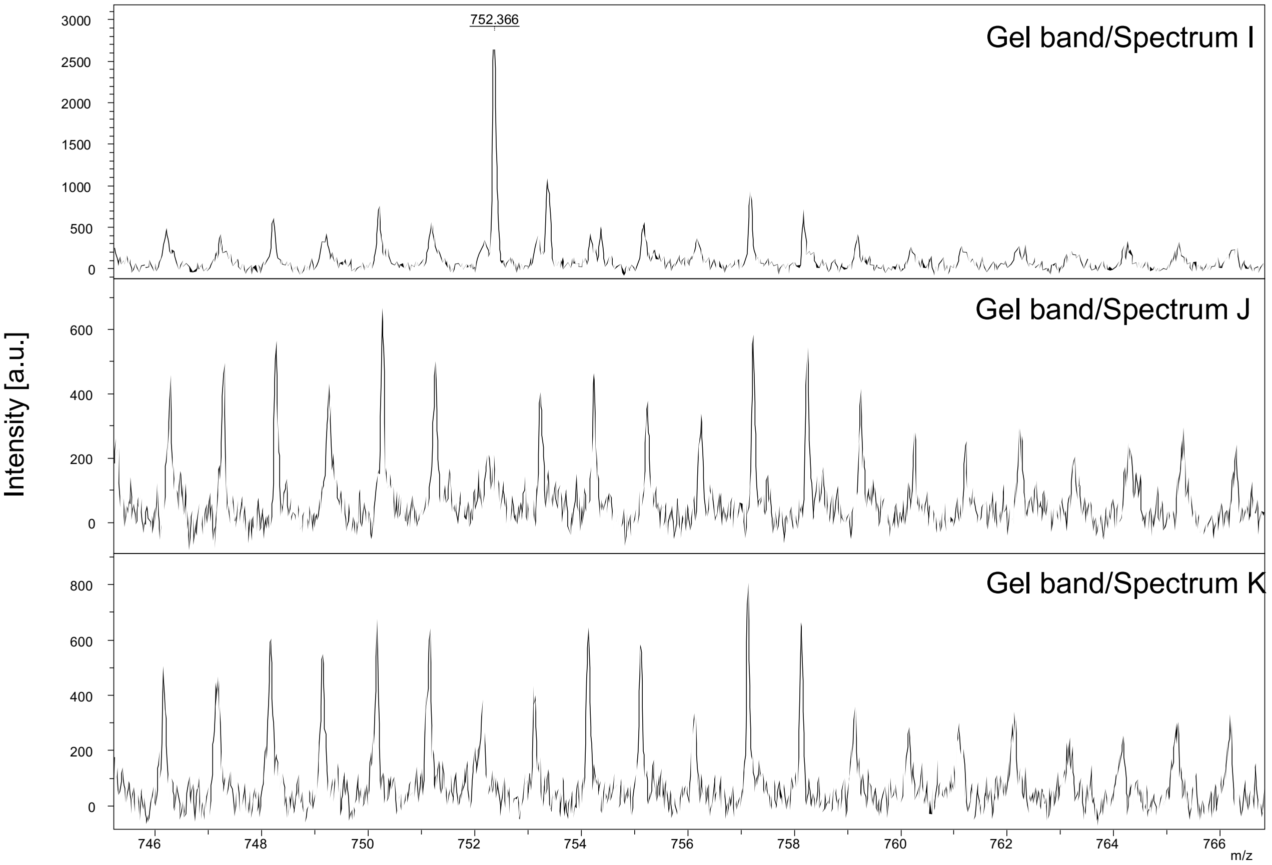

752.366
Gel band/Spectrum I
Gel band/Spectrum J
Gel band/Spectrum K
Intensity [a.u.]
m/z


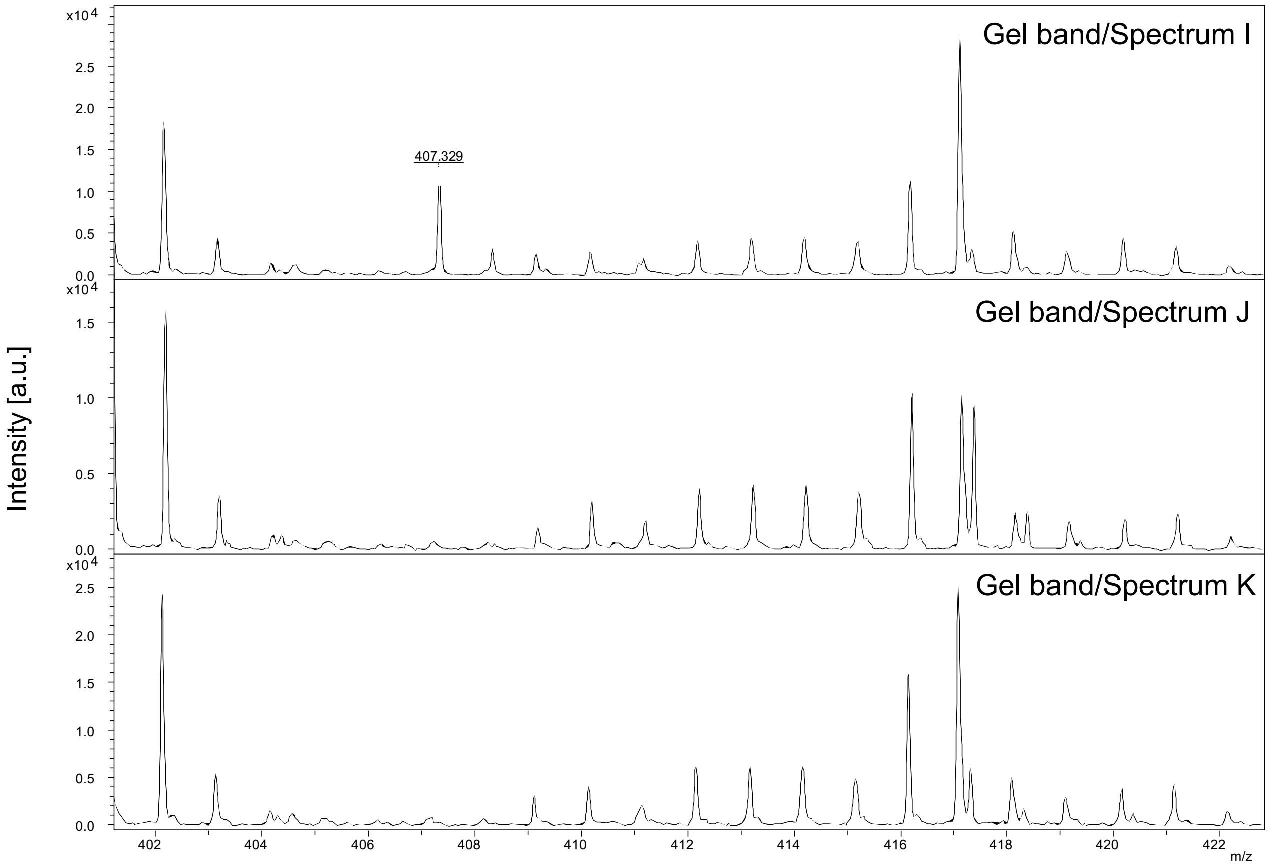

407.329
Gel band/Spectrum I
Gel band/Spectrum J
Gel band/Spectrum K
Intensity [a.u.]
x10^4
m/z

Gel band/ Spectra L, M and N (PSMα4)

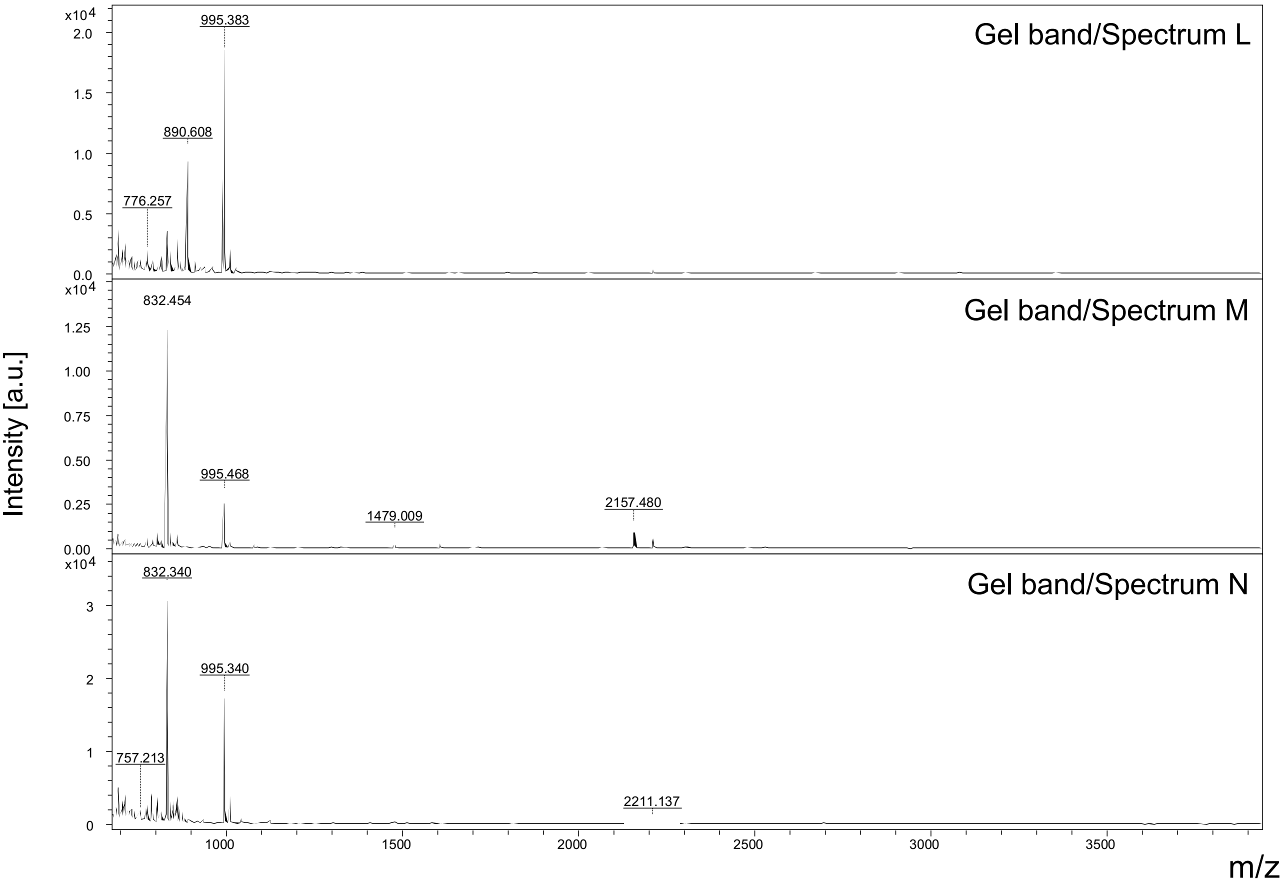


*MS spectra of in-gel digested PSMα4 samples*. **Gel band/Spectrum: L** (2 kDa band), sample with 77.7 μM PSMα4; **M** (37 kDa band) and **N** (120 kDa band) sample with pooled digests of 25 μM α-syn + 0.97 μM PSMα4 and 25 μM α-syn + 77.7 μM PSMα4. The following PSMα4 peptide was detected in gel band L and is presented in the zoomed spectrum below: AIIDIFAK (aa from 13 to 20) with a theoretical mass of 890.53 Da.

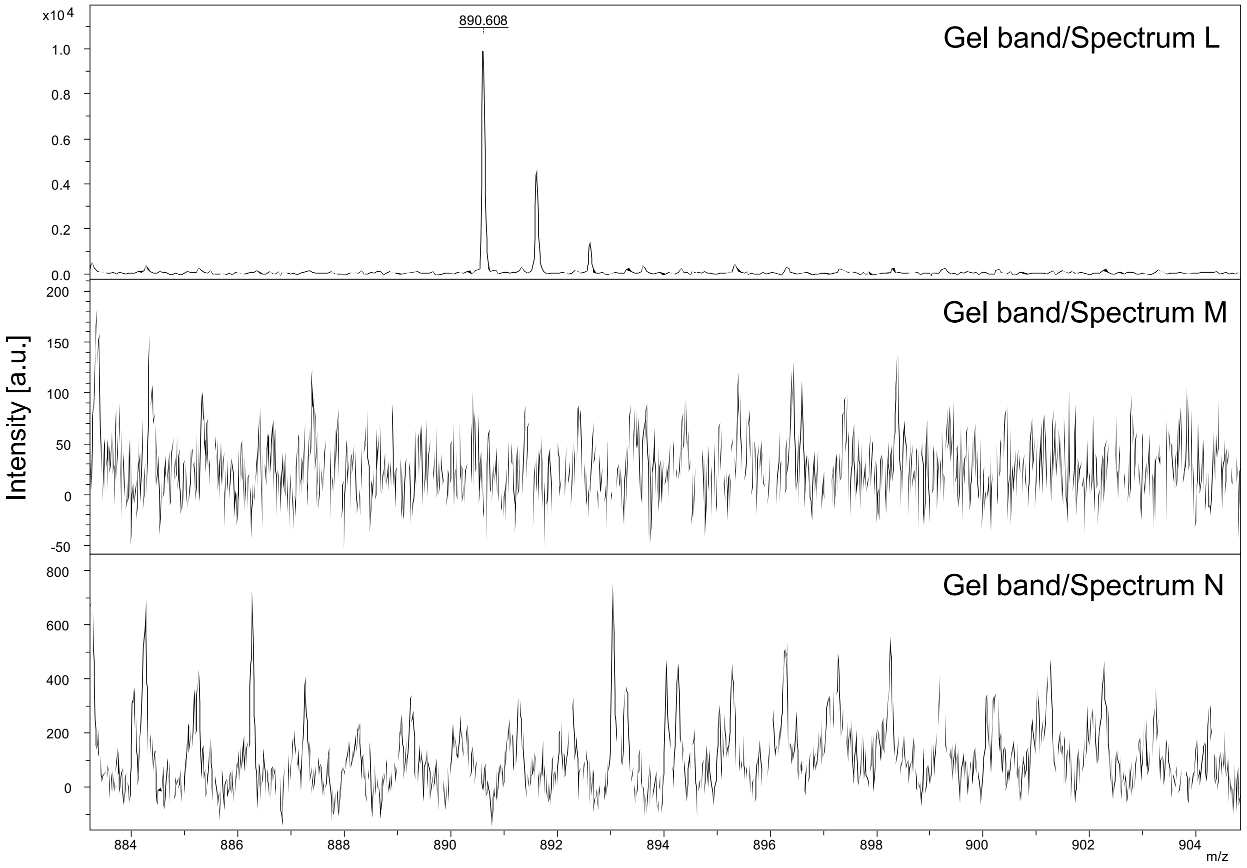


Supplementary Figure 9. α-syn and PSMα peptides do not form heteromolecular aggregates. A-syn and PSMα peptides incubated separately or together in Tris buffer were size separated on a Tricine SDS gel. The monomer α-syn bands, monomer PSMα peptide bands and the 37 kDa and 120 kDa bands were digested and analyzed by mass spectrometry. While the expected peptides of all the PSMα peptides were identified in the <10 kDa bands, only α-syn was detected in the 37 and 120 kDa bands.

asyn-GFP pSyn

7.8 µM PSMα1

2.5 µM α-syn (1.1% DSMO)

2.5 µM α-syn + 0.86 µM PSMα1

2.5 µM α-syn + 0.3 µM PSMα1

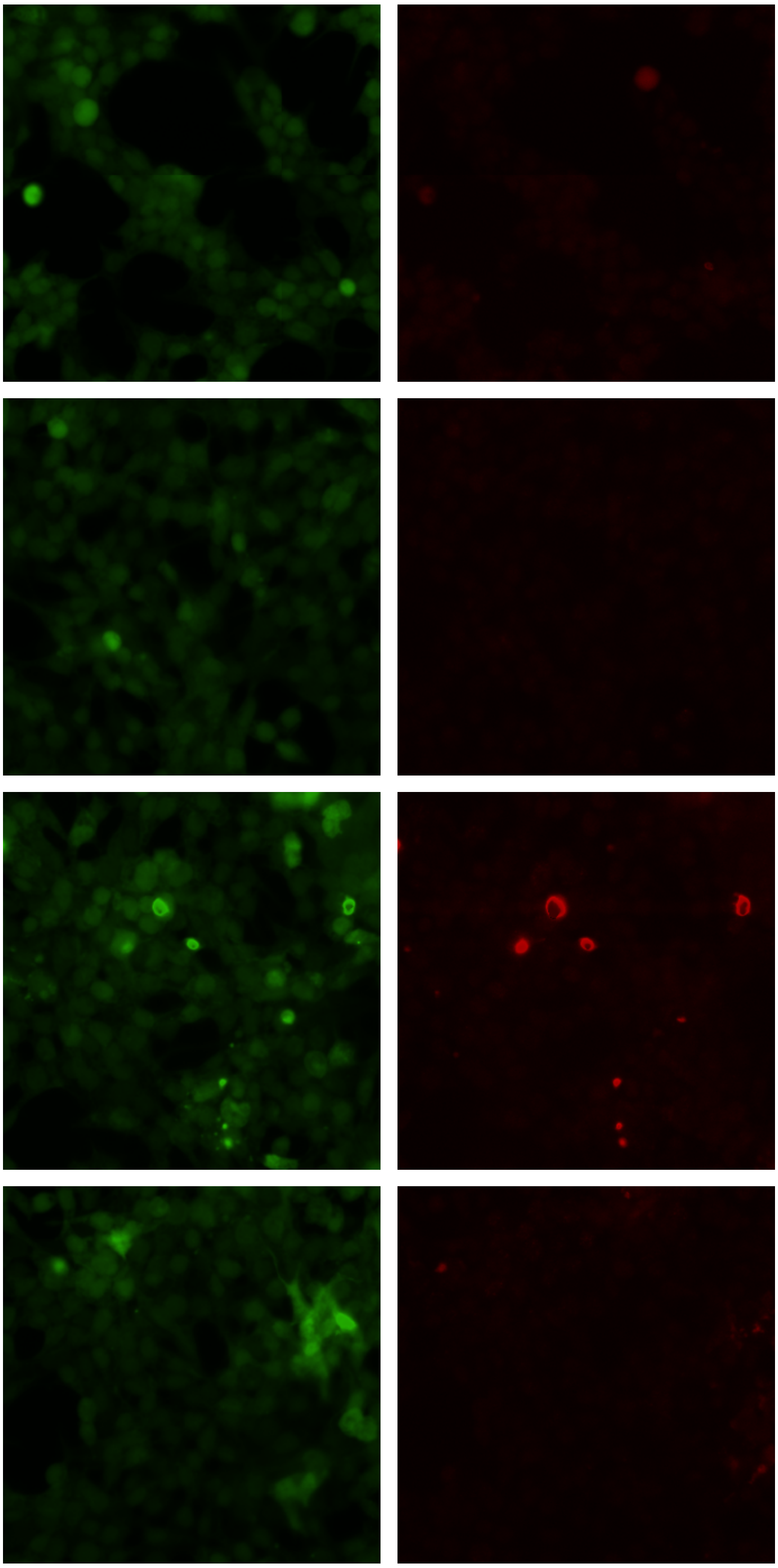

asyn-GFP pSyn

7.8 µM PSMα2

2.5 µM α-syn (1.1% DSMO)

2.5 µM α-syn + 2.6 µM PSMα2

2.5 µM α-syn + 0.86 µM PSMα2

2.5 µM α-syn + 0.3 µM PSMα2

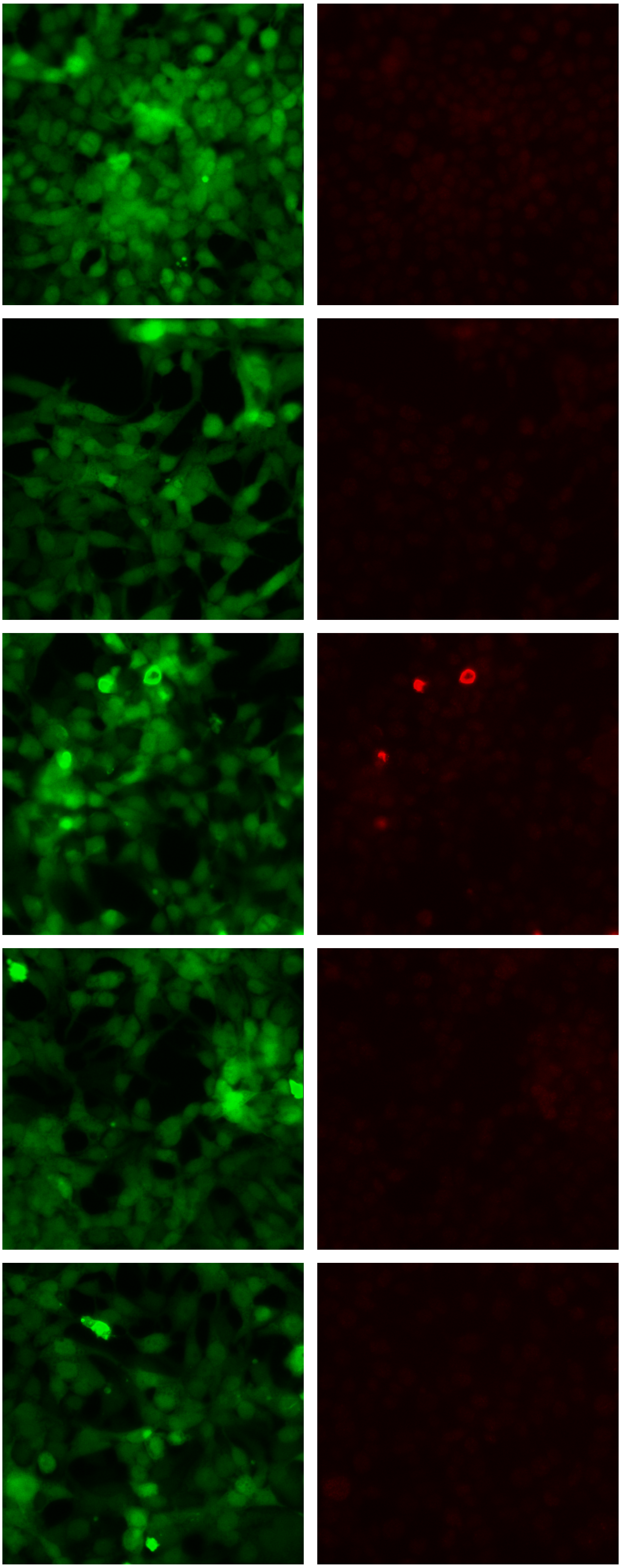

asyn-GFP pSyn

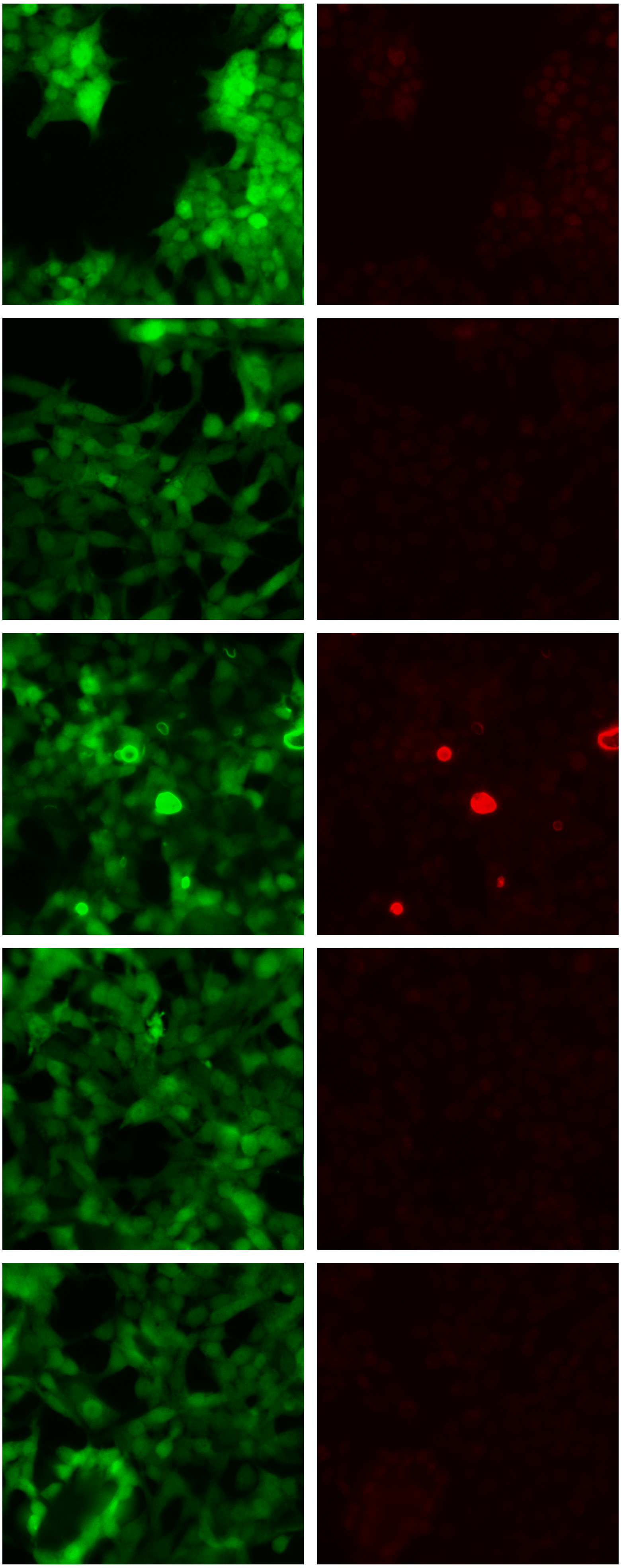

7.8 µM PSMα3

2.5 µM α-syn (1.1% DSMO)

2.5 µM α-syn + 2.6 µM PSMα3

2.5 µM α-syn + 0.86 µM PSMα3

2.5 µM α-syn + 0.3 µM PSMα3

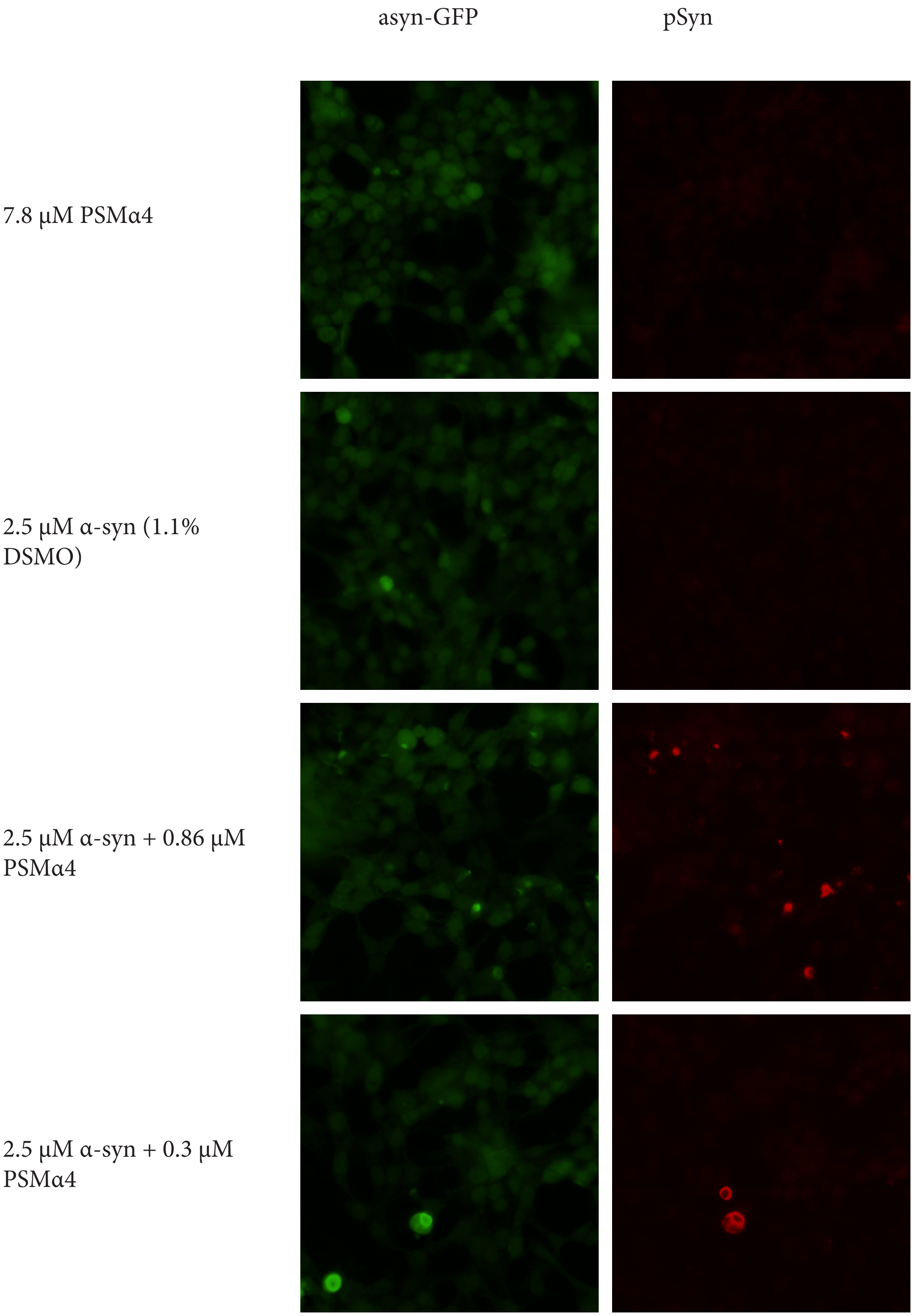
asyn-GFP
pSyn
7.8 µM PSMα4
2.5 µM α-syn (1.1% DSMO)
2.5 µM α-syn + 0.86 µM PSMα4
2.5 µM α-syn + 0.3 µM PSMα4

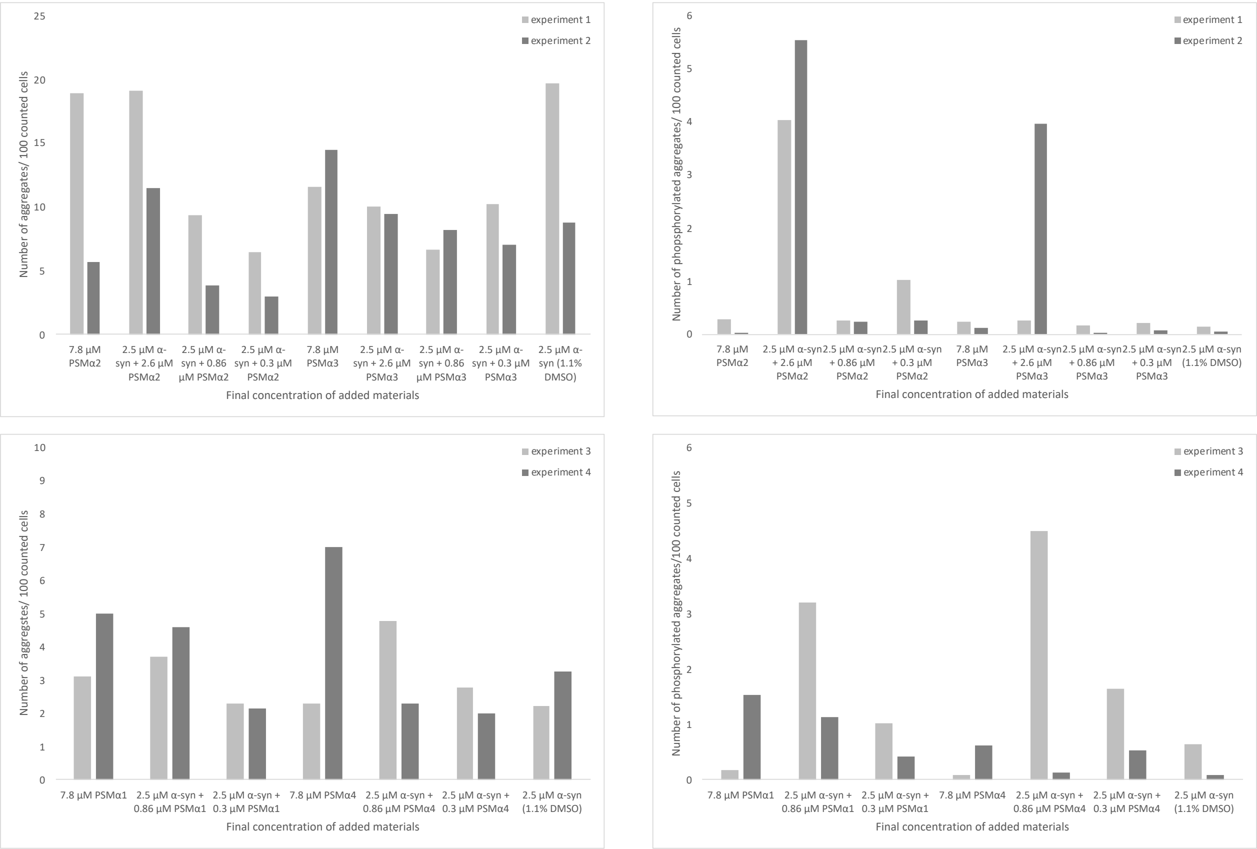


Supplementary Figure 10. α-syn aggregated in the presene of PSMα peptides induces seeding and phosphorylation in cells. HEK cells were stained for α-syn phosphorylated at ser129 after 48 hour-treatment with α-syn aggregated on its own or in the presence of PSMα peptides (final concentrations of added materials shown). Representative images from one well from each treatment condition are shown. These experiments were repeated twice. The number of aggregates and phsophorylated aggregates were quantified per 100 counted cells. The averages from all replicate wells are shown for each experiment. The representative images shown are from experiments 2 and 3.
